# IKAROS Gene Regulatory Network Reveal ERG as a Vulnerability in B-cell Acute Lymphoblastic Leukemia

**DOI:** 10.64898/2025.12.29.696402

**Authors:** Hana Paculova, Alyssa Richman, Joseph Boyd, Joshua Carlos Bernil, Princess Rodriguez, Etapong Ayongaba, Prachi Ghule, Elsa Drivet, Amrita Palaria, Mariam Dajani, Els Verhoeyen, Anthony Saviola, Gary S. Stein, Hilde Schjerven, Seth Frietze

## Abstract

B-cell acute lymphoblastic leukemia (B-ALL) is driven by transcriptional dysregulation that impairs B-cell differentiation and sustains leukemic growth. A defining feature of high-risk B-ALL is mutations in *IKZF1*, which encodes the tumor suppressor IKAROS. Here, we map IKAROS gene regulatory networks in *IKZF1*-mutated Ph B-ALL using an inducible IKAROS system and multi-omic profiling. IKAROS restoration reprograms chromatin accessibility and transcriptional control, shifting regulation from an ETS-dominated state to one enriched for B-cell lineage factors. Among repressed transcription factors, we identify ERG as a key regulatory node directly bound and antagonized by IKAROS. IKAROS binds regulatory elements near *ERG* and other progenitor-associated genes, coinciding with reduced *ERG* expression and repression of transcriptional programs linked to early B-cell developmental stages. Analysis of single-cell multiome data from human B-cell progenitors shows that ERG and IKAROS have opposing stage-specific activities and identifies a developmental stage-specific regulatory region in *ERG* intron 3 which is bound by IKAROS, and functionally important for *ERG* gene expression. Functional assays using CRISPRi and ETS inhibitors, along with gene dependency data from DepMap, confirm ERG dependency in *IKZF1*-deficient B-ALL. Our findings identify ERG as a context-specific dependency in *IKZF1*-deficient B-ALL, providing a mechanistic basis for the observed mitigation of poor prognosis for *IKZF1*-mutation in patients with co-occurring *ERG* deletions.

## Introduction

Acute lymphoblastic leukemia (ALL) is the most common childhood malignancy, with B-cell precursor ALL (B-ALL) accounting for most pediatric and adult cases [1]. B-ALL arises from malignant transformation of progenitor B-lineage lymphocytes in the bone marrow, leading to impaired hematopoiesis, uncontrolled cell growth and developmental arrest at the pro-B or early pre-B stage, prior to formation of a functional B-cell receptor [2, 3]. Although once considered a relatively uniform disease, transcriptional and genetic profiling have revealed extensive heterogeneity, with at least 14 subgroups defined by gene expression [4] and up to 28 variants recognized when morphologic, genetic, and clinical features are considered [5].

Large-scale genomic studies have shown that most B-ALL subtypes harbor alterations in genes encoding hematopoietic transcription factors (TFs) or their regulators, including *PAX5*, *ETV6*, *IKZF1*, and *RUNX1*, highlighting the central role of TF dysfunction in leukemogenesis [5–7]. These factors normally coordinate lineage-specific gene expression by shaping chromatin structure at *cis*-regulatory elements required for B-cell differentiation. Their disruption alters enhancer–promoter interactions, blocking normal differentiation and enabling persistent leukemic transcription [8–10]. Enhancer dysregulation is now recognized as a hallmark of hematologic malignancies, through enhancer reprogramming, hijacking, or *de novo* creation, and oncogenic TFs often maintain leukemic states through long-range enhancer–promoter interactions [11]. These dependencies are increasingly viewed as therapeutic vulnerabilities, with BET and CDK7/9 inhibitors showing promise in preclinical models [12].

Philadelphia chromosome-positive (Ph) B-ALL, defined by the oncogenic BCR-ABL1 fusion, is a high-risk subtype with distinct transcriptional features and poor prognosis, particularly in adults [13]. Despite clinical benefit from tyrosine kinase inhibitors, relapse is common, especially in cases with additional genetic lesions. Among these, mutations in *IKZF1*, encoding the C2H2 zinc finger TF IKAROS, are strongly associated with treatment resistance and adverse outcome [14–17]. *IKZF1*, located on chromosome 7p12.2, is recurrently altered in B-ALL through a variety of mechanisms. Copy number studies show *IKZF1* locus deletions in ∼15% of childhood B-ALL and 40–50% of adult cases [6, 7, 18, 19]. These include whole-gene deletions (Δ1–8) that abolish *IKZF1* expression, deletions spanning exons 2–3, 2–7, or 4–8 [20], as well as point mutations [21, 22]. Furthermore, a frequent RAG-mediated intragenic focal deletion generates the dominant-negative isoform IK6 (Δ4–7) [23]. Most B-ALL have monoallelic *IKZF1*-mutations, but biallelic lesions also occur in a subset of cases [21, 24].

Functionally, many of these lesions impair DNA binding while retaining protein–protein interaction domains, particularly the IKAROS dimerization domain encoded in the terminal exon 8, allowing mutant isoforms to sequester both wild type IKAROS and other *IKZF* family members and associated chromatin remodeling complexes. Clinically, *IKZF1* lesions defines the very high-risk “*IKZF1^plus^*” subgroup when combined with *CDKN2A*, *CDKN2B*, *PAR1*, or *PAX5* deletions, and in the absence of *ERG* deletion [25, 26].

IKAROS is a central regulator of B-lineage identity, chromatin structure, and gene expression during lymphoid development. It controls enhancer–promoter looping and chromatin accessibility at immune regulatory loci [27, 28]. While its critical role as a tumor suppressor function is established in both murine models and human patient data, the mechanisms by which IKAROS regulates enhancer architecture and transcriptional programs in human B-ALL remain less well understood. To investigate mechanisms of IKAROS-mediated tumor suppression in human B-ALL, we here used an inducible system to restore full-length IKAROS (IK1) in patient-derived Ph B-ALL cells harboring *IKZF1* lesions. We used this system to capture early transcriptional and chromatin changes following IKAROS expression, before the completion of growth arrest. IKAROS re-expression modified chromatin accessibility and transcription factor networks, recapitulating several regulatory changes that normally occur as progenitors progress toward B-cell commitment. A major effect of IKAROS restoration was suppression of the ETS-family transcription factor ERG, which is highly expressed in *IKZF1*-mutated B-ALL and normally downregulated during later stages of B-cell development. These findings suggest that B-ALL associated mutations in *IKZF1* disrupts developmental repression of ERG and related networks, and that IKAROS tumor suppressor function involves direct antagonism of early-developmental transcriptional circuits.

## Results

### IKAROS re-expression reconfigures the transcriptional regulatory network of *IKZF1*-mutated Ph□ B-ALL

Prior studies demonstrated that IKAROS expression in *IKZF1*-mutated Ph^+^ B-ALL cell lines induced cell cycle exit and suppressed leukemic growth [29, 30]. To investigate the underlying gene regulatory mechanisms, we reintroduced wild-type full-length IKAROS (IK1) using a doxycycline-inducible lentiviral system in two patient-derived *IKZF1*-mutated Ph B-ALL cell lines (Fig. 1A). These cell lines (MXP5 and PDX2) harbor heterozygous deletions spanning *IKZF1* exons 4–6, disrupting the N-terminal zinc-finger DNA-binding domain creating the dominant negative IK6 isoform. Both display features of an early B-cell precursor phenotype and serve as representative models of *IKZF1*-mutant Ph B-ALL [30]. To capture early regulatory changes preceding overt cell-cycle arrest, we performed RNA-seq and ATAC-seq 24 hours after IK1 induction, a time point at which cells begin to exit the cell cycle (Fig. S1A). PCA of RNA-seq and ATAC-seq revealed distinct baseline profiles for each cell line, with both gene expression and chromatin accessibility shifting similarly after IK1 induction (Fig. 1B, left).

**Figure 1.**
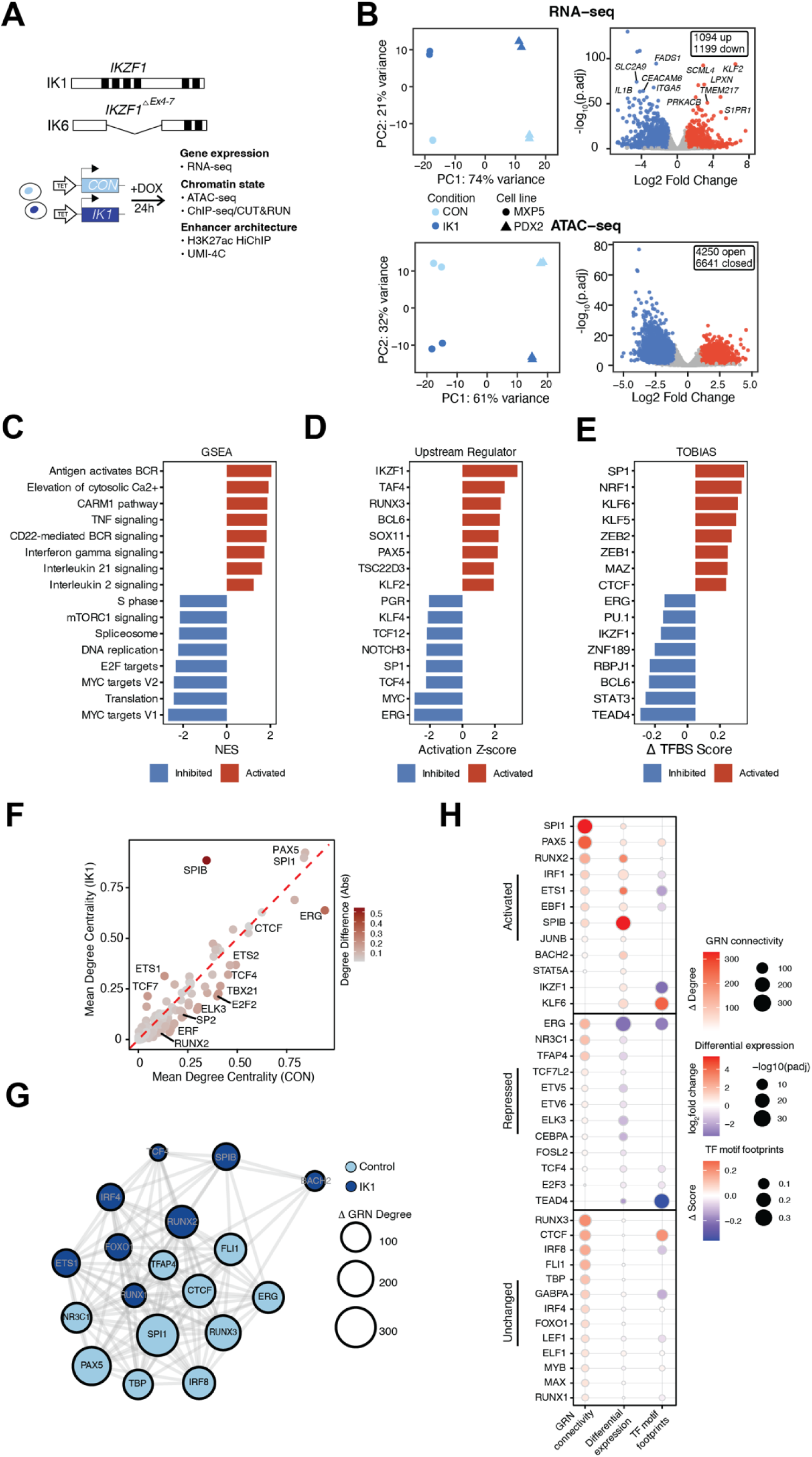
Induction of full-length Ikaros isoform IK1 changes gene expression and chromatin accessibility programs to rewire regulatory networks in Ph^+^ B-ALL cells. **A)** Schematic of the doxycycline-inducible IK1 expression system used in *IKZF1*-mutated Ph^+^ B-ALL cell lines. Cells expressing full-length *IKZF1* (Ik1-IRES-GFP; IK1) or empty vector control (GFP only; CON) were harvested 24 hours after DOX treatment for integrated analysis of gene expression, chromatin accessibility and epigenomic profiling using RNA-seq, ATAC-seq, ChIP-seq/CUT&RUN, HiChIP and UMI-4C. **B)** RNA-seq and ATAC-seq analyses of MXP5 and PDX2 cells following 24-hour IK1 induction. Left: Principal component analysis (PCA) of RNA-seq (top) and ATAC-seq data (bottom) showing separation by condition (IK1 vs. CON) and cell line (MXP5 vs. PDX2). Right: Volcano plots showing differentially expressed genes (DEGs) (top) and differentially accessible regions (DARs) (bottom). RNA-seq analysis identified 2,293 DEGs (adjusted p < 0.05, |log FC| > 1), including 1,094 upregulated and 1,199 downregulated genes. Selected DEGs are labeled. ATAC-seq analysis identified 10,891 DARs, with 4,250 regions gaining accessibility (open) and 6,641 showing reduced accessibility (closed) following IK1 induction (adjusted p < 0.05, |log FC| > 1). **C)** Gene set enrichment analysis of IK1-responsive genes with canonical pathway enrichment analysis of differentially expressed genes (DEGs) using MSigDB and curated gene sets. **D)** Upstream regulator analysis performed using Ingenuity Pathway Analysis (IPA). Shown are TFs with the top activation Z-scores (FDR < 0.05). **E)** Differential TF footprint score analysis based on ATAC-seq data (TOBIAS) under IK1 and control conditions. Shown are transcription factor motifs with significantly increased (activated; red) or decreased (inhibited; blue) corrected footprint scores following IK1 induction. **F)** Comparison of mean GRN degree centrality of TFs in control versus IK1. Each point represents a TF, with position indicating mean GRN degree (i.e., connectivity to target genes) under control (x-axis) and IK1 (y-axis). TFs above the diagonal (red dashed line) show increased connectivity in IK1, while those below exhibited increased target gene connectivity in CON. Color scale based on absolute degree difference (|Δ degree|). **G)** TF regulatory network derived from IK1-responsive gene regulatory changes. Nodes represent TFs changes in network connectivity (degree) between control and IK1. Node color indicates the condition where TF exhibits largest regulatory activity (light blue = control, dark blue = IK1), and node size is scaled to the absolute change in GRN degree (|Δ degree|). Edges represent shared target genes (TGs) between TFs, with edge thickness proportional to the number of co-regulated targets. **H)** Summary of IK1-responsive TFs grouped by RNA expression changes (Activated, Repressed, Unchanged). Each row is a TF. Columns show: GRN connectivity, where point size and red intensity represent the magnitude of change in connectivity (|Δ degree|, IK1 vs. control) without distinguishing direction; Differential expression, where color indicates log fold change (red = up, blue = down in IK1) and point size encodes significance (–log adjusted p); and TF footprinting, where color shows direction of footprinting change (red = increased, blue = decreased binding in IK1) and point size reflects |Δ score|.

Differential expression analysis identified 2,293 genes significantly regulated by IK1 across both cell lines (|log FC| > 1, FDR < 0.05) (Fig. 1B, right, top, Table S1). Concurrently, 10,891 differential accessibility regions (DARs) were observed, with 61% (6,641) decreasing and 39% (4,250) increasing in accessibility, consistent with a dual role for IKAROS in chromatin remodeling through both compaction and opening (Fig. 1B, right, bottom, Table S2). Gene set enrichment analysis showed IK1-activated genes involved in B-cell receptor signaling, calcium signaling, and immune-related regulatory networks (Fig. 1C, Fig. S1B, Table S3). Conversely, downregulated genes were enriched for MYC targets, E2F targets, mTOR signaling, cell cycle and DNA replication, consistent with the induction of growth arrest. To define the transcriptional programs responsive to IK1, we integrated TF activity inference, TF footprinting, and gene regulatory network analysis. Upstream regulator analysis (Ingenuity Pathway Analysis) identified IKAROS as the top activated TF, consistent with its re-expression (Fig. 1D; Table S4). Other predicted activators included PAX5, BCL6 and KLF2, while repressed regulators included MYC, ERG, and SP1. To further assess TF motif occupancy changes, we used TOBIAS (Transcription factor Occupancy prediction By Investigation of ATAC-seq Signal) footprinting analysis on control and IK1-induced ATAC-seq profiles [31]. IKAROS motifs showed significantly reduced accessibility following IK1 induction, consistent with a chromatin-compacting role (Fig. 1E, S Fig. 1C, Table S5). Footprint for ERG, TEAD, PU.1, BCL6 and STAT3 also showed decreased occupancy, many of which are associated with early hematopoietic or leukemic programs. In contrast, increased occupancy was observed at KLF6, MAZ, SP1, and ZEB2 motifs.

To evaluate how IK1 re-expression influences the transcriptional network structure, we constructed gene regulatory networks (GRNs) under control and IK1 conditions by integrating RNA-seq and ATAC-seq data. TFs were linked to candidate target genes based on expression correlation and the presence of TF motifs within accessible *cis*-regulatory elements associated with those genes [32]. Comparison of these networks revealed condition-specific shifts in TF connectivity to target genes. TFs such as SPI1, PAX5, and SPIB exhibited increased connectivity in the IK1 network, whereas factors including ERG, E2F2 and ETS2 lost predicted regulatory interactions upon IK1 induction (Fig. 1F, Fig. S1D,E, S Table S6). IK1 induction led to extensive rewiring of transcriptional circuitry, with TF–TF network analysis indicating redistribution of regulatory influence among B-cell transcription factors (Fig. 1G). TFs were classified by whether they showed greater connectivity in the IK1 or control network. The relative change in connectivity is represented by node size, with larger nodes indicating TFs whose degree shifted most strongly between conditions. For example, RUNX2 gained interactions in the IK1 network, while ERG and SPI1 were more highly connected in the control network. These changes could reflect shifts in TF expression, binding activity, or both. To distinguish these mechanisms, TFs were categorized based on their expression changes and grouped into activated, repressed, or unchanged in response to IK1. Activated TFs (SPI1, PAX5, RUNX2) showed increased RNA expression following IK1 induction, with high GRN connectivity, and PAX5 also had enhanced motif accessibility (footprint), suggesting that IK1 promotes their regulatory activity (Fig. 1H). In contrast, repressed TFs including ERG, SPI1, PAX5, RUNX3, and FLI1 exhibited reduced GRN connectivity following IK1 induction. Among these, ERG showed the largest decrease in GRN connectivity (Fig. 1F), supported by reduced expression, diminished footprint scores, and predicted downregulation of ERG target genes (Fig. 1D–E). CTCF, RUNX3, and IRF8 maintained stable expression but displayed changes in GRN connectivity (Fig. 1H). Among these, IRF8 also exhibited reduced motif accessibility, consistent with its known role in immune regulation and the observed downregulation of immune-related pathways following IK1 induction (Fig. 1C, S Table 1). To evaluate whether the TFs identified to be changed between the control vs IK1-induced GRN reflects defining regulatory factors in *IKZF1*-mutated B-ALL, we analyzed RNA-seq and ATAC-seq data from a panel of *IKZF1*-wild-type (WT) and *IKZF1*-mutated (IK6) B-ALL cell lines (Fig. S1F). GRN analysis of these two groups revealed differences in TF connectivity, with 56.3% of TF–target interactions shared between groups and the remainder unique to either IK6 or WT networks (Fig. S1G). Notably, IK6 networks more closely resembled the pre-induction (CON) state observed in the IK1 model, whereas the WT group network aligned with the IK1-induced condition. These data show that TF regulatory networks in *IKZF1*-mutant B-ALL cell lines more closely resemble the control condition prior to IK1 induction, while WT cell lines correspond to the IK1-induced network state. This pattern supports a role for IKAROS influence on TF activity in B-ALL cells, where reduced IKAROS activity in mutant B-ALL cells correlate with higher ERG connectivity, which is diminished following IK1 induction. Conversely, IK1 induction reduces ERG-centered network activity while increasing SPIB regulatory activity (Fig. S1F).

### Ikaros re-expression establishes transcriptional changes consistent with B-cell maturation

Since IK1 induction altered TF activity in Ph B-ALL, we next asked whether these changes correspond to normal developmental programs in B-cell differentiation. We analyzed paired single-cell RNA-seq and ATAC-seq (10X multiome) data integrated from ten healthy bone marrow donors. Among diverse hematopoietic populations, we identified and isolated B-cell lineage precursors, characterized by stage-specific gene expression and chromatin accessibility patterns across both modalities (Fig. 2A, Fig. S2). These included hematopoietic stem cells (HSCs), common lymphoid progenitors (CLPs), and successive stages of B-cell precursors (pro-B, pre-B, and immature B cells). These populations trace the developmental progression from multipotent progenitors to lineage-committed B cells. Early stages (HSCs and CLPs) showed enrichment for stem and lymphoid genes (*CD34*, *KIT*, and *FLT3*), while later B-cell precursor stages expressed B cell identity factors (*VPREB1*, *IGLL1*, *MS4A1*, and *HLA-DQA1*) (S Fig. 2). IK1-induced GRN TFs followed stage-specific expression patterns. Activated TFs, including *EBF1*, *BACH2*, *ETS1*, *SPIB*, *IRF1* were upregulated during the pro-B to immature B-cell stages. *RARA* and *IKZF1* were relatively consistently expressed across each stage with *IKZF1* expression increasing most in the immature B cell stages. Repressed TFs, including *ERG, BCL11A, ETV6, ELK3, NR3C1* and *FOXO3*, showed highest expression in early progenitors that declined with maturation following proB stage. The unchanged group maintained relatively stable expression across stages. When scored across single cells, repressed TFs were most active in progenitors, whereas activated TFs rose during the pro-B to immature B-cell stages, and unchanged TFs showed little variation (Fig. 2C,D). We then inferred TF regulatory activity using SCENIC+, which integrates motif accessibility and target gene co-expression [33]. Early progenitors were enriched for ETS/hematopoietic regulons (ERG, GATA2, C/EBP), whereas later stages showed induction of B-lineage regulons (PAX5, IRF4, IKZF family) (Fig. 2E). In SCENIC+, signs denote the direction of change for gene-based (targets) and region-based (motif accessibility) activity; +/+ indicates coordinated activation, −/− coordinated repression, and −/+ reduced target expression despite higher motif accessibility, a pattern consistent with direct binding of a repressor or context-dependent repression. Because position weight matrices (PWMs) often overlap within TF families, SCENIC+ reports extended regulons (eRegulons), which represent the collective activity of TFs with shared binding motifs and predicted targets. We aligned activity trajectories of these eRegulons with stage-resolved RNA expression and assigned representative gene-level labels. Based on expression trends for *ERG* and *IKZF1* across differentiation, we identified two IKZF1 eRegulons, one repressive (−/−) and one with reduced activity (−/+), as well as an activating ERG eRegulon (+/+). The ERG_extended +/+ regulon showed peak activity in early progenitors and declined after the pro-B stage, whereas IKZF1_extended −/− activity increased during later stages, particularly from pre-B to immature B cells (Fig. 2E-G). These changes in regulon activity mirror the pseudotime trajectories of ERG and IKZF1 RNA expression, consistent with a developmental transition from ERG-driven progenitor programs to IKZF1-associated repression during B-cell maturation (Fig. 2G,F).

**Figure 2.**
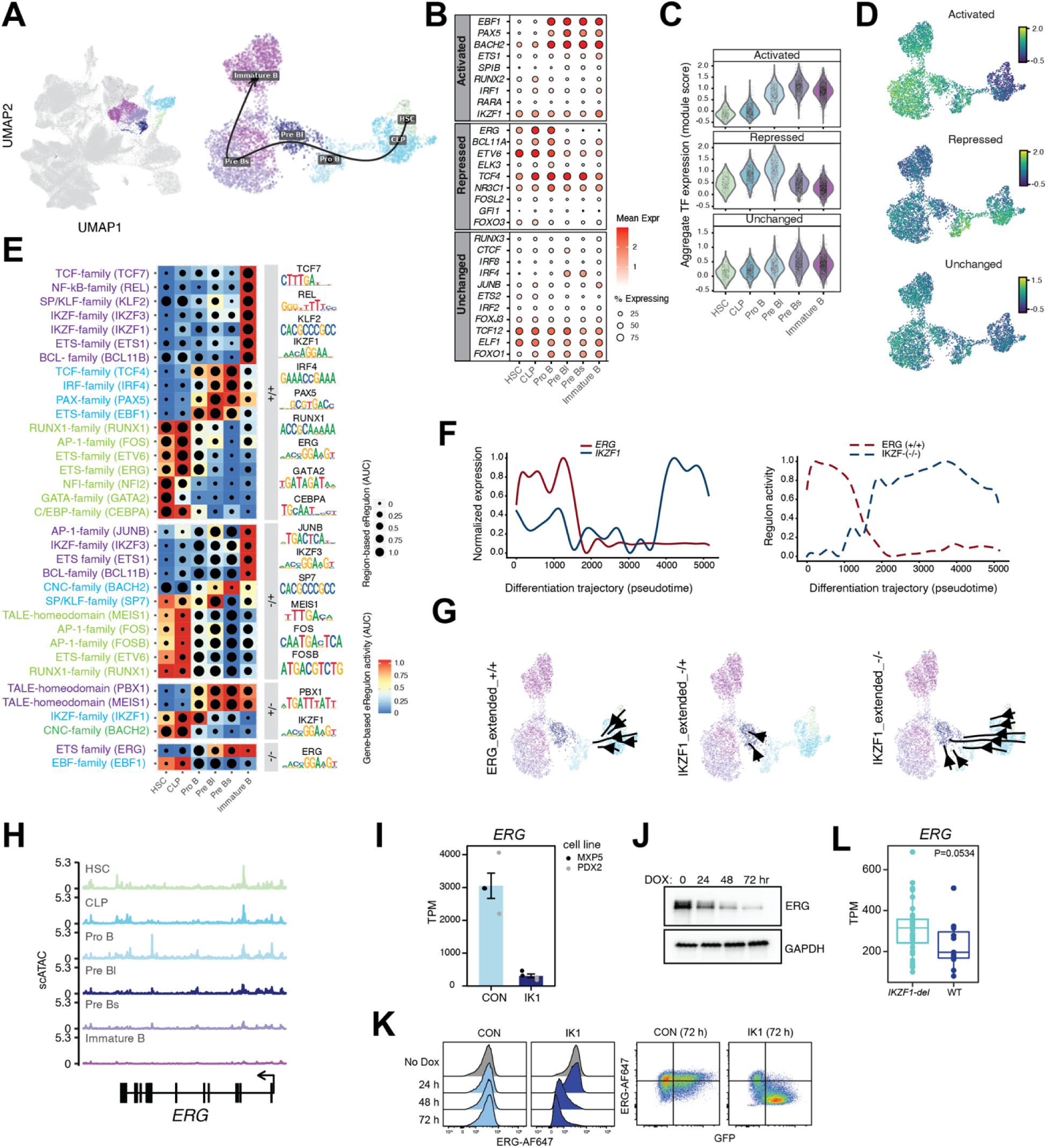
IKAROS-regulated transcription factor networks parallel stage-specific regulation in normal B-cell development. **A)** UMAP of the RNA modality from 10x Genomics Multiome single cells, integrated across 10 healthy bone-marrow donors (GSE194122). Non-B lineages are shown in gray; B-cell precursor subsets are labeled. Arrow indicates developmental progression from HSC to Immature B. **B)** Expression of IK1-responsive transcription factors defined in MXP5 cells (Activated, Repressed, Unchanged) across healthy B-cell precursor stages; point size corresponds to the fraction of cells expressing gene and the color corresponds to the mean expression levels. **C)** Module scores for IK1-responsive TF sets (Activated, Repressed, Unchanged; same TFs as in B) across B-cell precursor stages. Higher scores indicate greater aggregate expression. **D)** UMAPs showing the distribution of the module scores from C) Activated, Repressed, and Unchanged TF groups across the B-cell precursor embedding. The color denotes score magnitude higher corresponds to greater aggregate expression level. **E)** SCENIC+ extended eRegulon activity across B-cell precursor stages. Rows show eRegulons annotated at the TF-family level based on PWM similarity (a representative TF is listed in parentheses). Heatmap color indicates gene-based activity (AUC of target-gene enrichment per cell, where AUC refers to the area under the recovery curve used to quantify regulon activity based on gene expression), and dot size indicates region-based activity (AUC of motif accessibility at predicted binding sites, derived from ATAC-seq). Representative PWM logos are shown on the right. Signs denote the direction of activity: +/+ indicates that both target-gene expression and motif accessibility increase; −/− indicates that both decrease; +/− indicates higher gene expression with lower motif accessibility; −/+ indicates lower gene expression with higher motif accessibility. **F)** Pseudotime trajectories of *ERG* and *IKZF1* expression and regulon activity. Left: smoothed RNA expression of *ERG* (red) and *IKZF1* (blue) across the B-cell differentiation pseudotime. Right: regulon activity scores for ERG (+/+) and IKZF1 (−/−) modules along the same pseudotime. **G)** Developmental UMAPs showing eRegulon activity gradients. Arrows indicate direction of increasing activity for ERG (+/+), IKZF1 (−/+), and IKZF1 (−/−) extended regulons across the B-cell precursor manifold. **H)** Pseudo-bulk scATAC-seq signal tracks at the *ERG* locus across B-cell precursor stages. Each track represents aggregated accessibility profiles from the indicated stage. Y-axis shows normalized read depth (RPM). **I)** Bulk RNA-seq expression of *ERG* (TPM) in Ph B-ALL cell lines (MXP5 and PDX2) under control (CON) or IK1 induction. Bars represent mean ± SEM across replicates; individual points correspond to each replicate. **J)** Immunoblot of ERG protein in MXP5 cells at 0, 24, 48, and 72 hours after doxycycline-induced IK1 expression. GAPDH is shown as a loading control. **K)** Flow cytometry analysis of intracellular ERG protein levels in PDX2 cells. Left: histograms of ERG-AF647 signal at baseline and at 24, 48, and 72 hours following IK1 induction, compared to control. Control is gated on live single cells. Dox-induced cells are gated on GFP^+^ live single cells. Right: representative gating of ERG-AF647 signal versus GFP at 72 hours after doxycycline-induced IK1 expression. **L)** RNA-seq expression of *ERG* (TPM) in primary Ph B-ALL samples from cohort EGAS00001007167 [34], stratified by *IKZF1* deletion status. Boxplots show median and interquartile range, P-value was calculated using a two-sided Wilcoxon rank-sum test.

Consistent with the developmental regulation of *ERG* expression, single-cell ATAC-seq tracks across normal B-cell precursors showed that chromatin accessibility at the *ERG* locus was highest in progenitors and diminished as cells matured (Fig. 2H). In Ph B-ALL cells, IK1 induction sharply reduced *ERG* mRNA levels in both MXP5 and PDX2 lines (Fig. 2I), accompanied by a time-dependent decrease in ERG protein by immunoblot (Fig. 2J). Flow cytometry confirmed loss of ERG protein at the single-cell level, with reduced signal evident by 48–72 hours after IK1 induction (Fig. 2K). Finally, analysis of primary Ph B-ALL samples showed that ERG expression was higher in cases harboring IKZF1 deletions compared to IKZF1 wild-type, suggesting that *IKZF1* deletion is associated with elevated *ERG* expression in primary B-ALL samples (Fig. 2L). These findings, together with developmental data, support a model in which IKAROS normally represses ERG as B cells mature, and IK1 induction in B-ALL re-establishes this regulatory program that is lost with *IKZF1* deletion.

### IKAROS re-expression disrupts ERG chromatin engagement through remodeling of shared regulatory elements

Re-expression of IKAROS in B-ALL perturbs ERG-associated transcriptional programs, linking its developmental role to tumor suppressor activity in leukemia. To explore this mechanism, we performed ChIP-seq for ERG and IKAROS under control and IK1-induced conditions and called high-confidence peaks from two biological replicates per condition in MXP5 cells. We identified 1,228 IKAROS peaks in control cells and 2,253 peaks following IK1 induction, with 881 peaks shared between conditions (∼72% of control peaks; ∼39% of IK1 peaks) (S. Fig. 3, Fig. 3A, Table S7). The lower number of peaks in control cells is consistent with reduced chromatin occupancy by endogenous IKAROS in *IKZF1*-haploinsufficient MXP5 cells, whereas IK1 induction resulted in 1,372 new IKAROS binding sites. In parallel, ERG occupancy decreased substantially with IK1 (2,258 control vs 1,262 IK1 peaks; ∼ 44% loss), consistent with its down regulation at 24 hours. Most ERG peaks in IK1 cells overlapped with those in control cells, indicating a contraction of the ERG cistrome rather than redistribution. To better understand how IK1 expression alters ERG and IKAROS binding, we stratified ChIP-seq peaks into groups based on condition-specific and shared occupancy: IKAROS CON-specific, IKAROS IK1-specific, IKAROS shared, ERG CON-specific, ERG IK1-specific, and ERG shared (referred to as clusters C1–C6; Fig. 3A, bottom). Clustering of ChIP-seq and ATAC-seq signals at each group revealed clear patterns of IKAROS–ERG co-occupancy in control cells and changes in chromatin binding upon IK1 induction (Fig. 3B). Notably, IKAROS CON-specific peaks (C1) exhibited strong IKAROS binding under control conditions that were lost with IK1, along with high ATAC-seq signal that diminished upon IK1 expression. IKAROS CON-specific peaks were associated with predominantly closed DARs and down-regulated DEGs (191 closed DARs and 86 downregulated DEGs vs. just no open DARs and 36 upregulated DEGs), consistent with loss of IKAROS binding and chromatin closure at these elements. IKAROS shared sites (C3) also showed reduced accessibility upon IK1 induction (109 closed, 2 open DARs) but were associated with both up- and down-regulated DEGs (28 and 55, respectively), indicating that sustained IKAROS binding results divergent regulatory effects. In contrast, ERG IK1-specfic sites (155 peaks; C5) showed increased IKAROS occupancy in IK1-induced cells and were exclusively associated with open DARs and mostly upregulated DEGs (12 open DARs and 26 upregulated DEGs), suggesting potential cooperation between IKAROS and ERG in gene activation at these loci. ERG-shared sites (C6) also gained IKAROS binding with IK1 induction and showed evidence of both activation and repression, with modest DAR closure and a mix of up- and downregulated DEGs (10 vs. 41 open and closed DARs; 128 up- vs. 86 down DEGs), consistent with dual regulatory roles at shared loci. Analysis of genomic binding distributions showed that IKAROS binding under control conditions was more frequently located in distal intergenic regions, whereas both IKAROS IK1 and ERG peak sets had a higher percentage of promoter-proximal sites (± 3kb from TSS) (Fig. 3C). Of 996 ERG peaks lost following IK1 induction, 214 (∼21.5%) became newly bound by IKAROS, with 38.8% of these located at promoter-proximal regions (±3 kb from TSS), suggesting selective redistribution to previously ERG-occupied elements. Motif analysis confirmed enrichment of the canonical ERG (GGAA/T) and IKZF1 (TGGGA) motifs within their respective peak sets (Fig. 3D). Although both motifs share short purine-rich cores, they belong to distinct TF families with divergent binding preferences. Clustering enriched motifs based on similarity of their position probability matrices grouped related sequences into transcription factor families (Fig. 3E). This revealed that ETS motifs were most strongly enriched in ERG-bound regions, particularly ERG CON-specific peaks, whereas IKAROS-associated regions showed higher enrichment for IRF and STAT motifs, consistent with known IKAROS co-regulatory interactions [35]. In contrast, ERG-associated and shared sites were enriched for AP-1 and NFκB motifs, aligning with the engagement of stress- and inflammation-related pathways observed upon IK1 induction (Fig. 1C).

**Figure 3.**
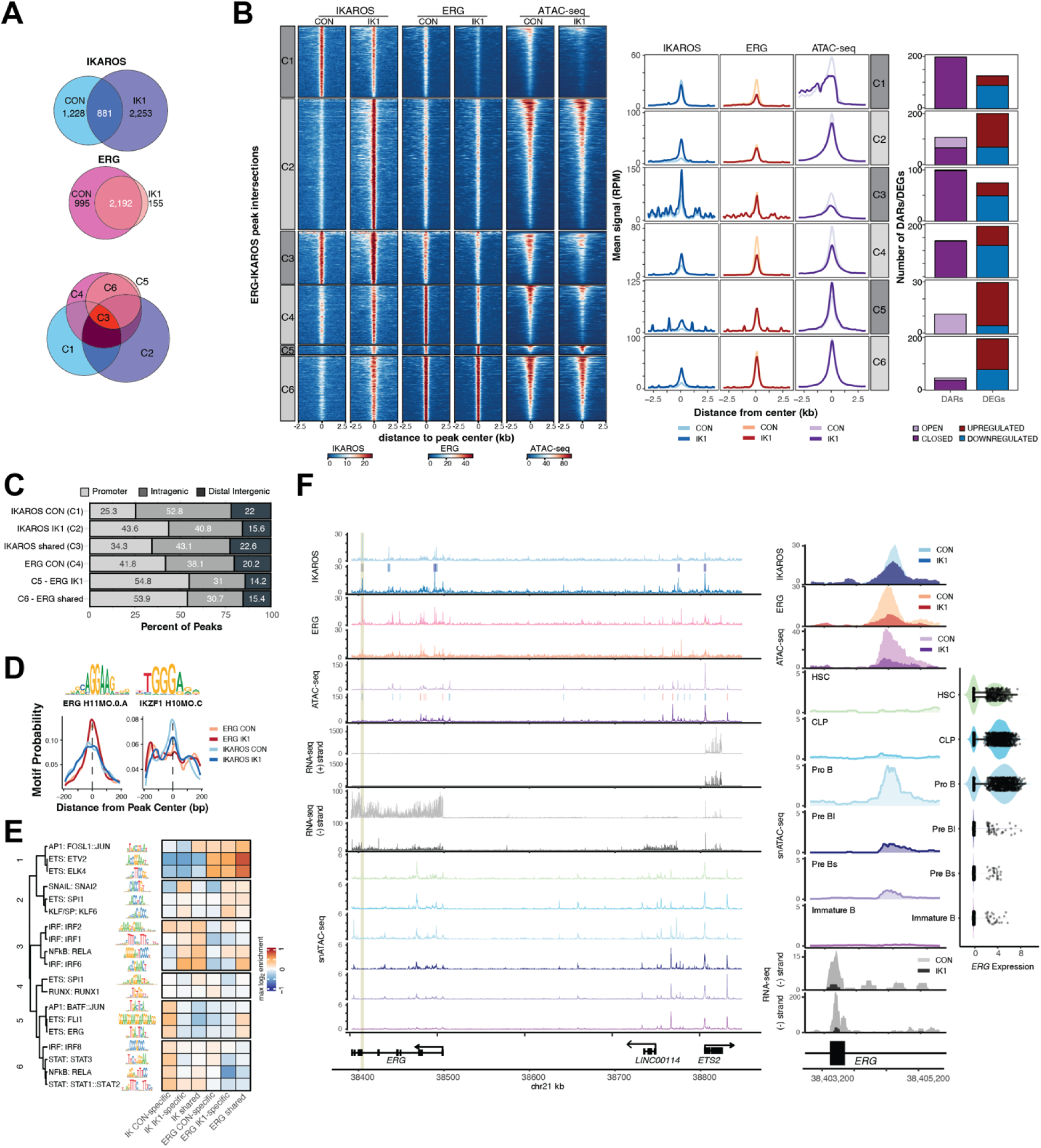
IKAROS and ERG binding dynamics reveal chromatin remodeling and direct regulation of *ERG* locus in Ph□ **B-ALL.** **A)** Overlap of ChIP-seq peaks for IKAROS and ERG in MXP5 cells under control (CON) and IK1-induced (IK1) conditions, with venn diagrams showing shared, CON-specific, and IK1-specific peak sets. Diagram below shows the shared and condition-specific intersections across all four peak sets. **B)** ChIP-seq peaks were stratified into six groups (IKAROS CON-specific, IKAROS IK1-specific, IKAROS shared, ERG CON-specific, ERG IK1-specific, and ERG shared). Heatmaps (left) show normalized signal intensity (RPKM) for IKAROS, ERG, and ATAC-seq within ±2.5 kb of peak centers across control and IK1-induced conditions, with corresponding average signal profiles shown on the right. Barplots summarize the number of open and closed DARs, as well as up- and down-regulated DEGs, associated with each peak group. Open DARs correspond to regions with increased ATAC-seq signal in IK1 compared to CON, while closed DARs show reduced accessibility in IK1, based on differential ATAC-seq analysis (|log FC| > 1, FDR < 0.05). **C)** Genomic distribution of IKAROS and ERG ChIP-seq peaks. Peaks were annotated as promoter-proximal (±3 kb from the transcription start site), intragenic (within gene bodies outside the promoter region), or distal intergenic (all other regions). Bar plots show the fraction of peaks in each category for control (CON), IK1-induced (IK1), and shared ERG and IKAROS peak subsets. Total peaks per group: IKAROS CON (n = 1207), IKAROS IK1 (n = 2211), IKAROS shared (n = 901), ERG CON (n = 996), ERG IK1 (n = 155), ERG shared (n = 1098). **D)** Motif enrichment profiles for canonical ERG and IKZF1 binding motifs. Sequence logos (top) show ChIP-seq derived (HOCOMOCO) consensus motifs for ERG and IKZF1 [36]. Line plots display motif probability across ±200 bp from the ChIP-seq peak center in ERG or IKAROS peak sets for control and IK1-induced conditions. Vertical dashed lines indicate the peak summit. **E)** Motif family enrichment across peak groups. Representative motifs are shown for enriched TF families, with sequence logos displayed to the left of the heatmap. Motifs were clustered by similarity of their position weight matrices into six higher-order clusters. The heatmap displays relative enrichment of these motif families across IKAROS and ERG ChIP-seq peak groups. ChIP-seq, ATAC-seq, RNA-seq, and single-cell ATAC-seq profiles across a ∼440 kb region of chromosome 21 spanning *ERG*, *LINC00114*, and *ETS2*. Tracks display IKAROS (light blue, CON; dark blue, IK1) and ERG (pink, CON; orange, IK1) ChIP-seq, ATAC-seq (purple, CON; dark purple, IK1), strand-specific RNA-seq (gray), and single-cell ATAC-seq across hematopoietic and B-cell stages. Significant ChIP-seq peaks and DARs are indicated above their respective tracks. The yellow vertical line marks an intronic IKAROS-bound region at ERG shown in the zoomed view (right), and violin plot of single-cell *ERG* expression.

Examination of a ∼440 kb region spanning the *ERG* locus identified multiple IKAROS binding sites, including control-specific, IK1-induced, and shared peaks that overlapped with regions of IK1-induced chromatin closing (Fig. 3F). A focal IKAROS binding site within an *ERG* intron coincided with reduced accessibility and downregulation of *ERG* expression following IK1 induction (Fig. 3F, right). While nearby *ETS2* expression remained unchanged, the adjacent long noncoding RNA *LINC00114* was upregulated with IK1, consistent with transcriptional remodeling across the locus. Developmental single-cell accessibility profiles revealed that the IK1-responsive intronic region becomes accessible at the Pro-B stage but is closed in early progenitors (HSC/CLP), whereas an upstream region near the *ERG* TSS is accessible in HSC/CLP and closes during B-cell commitment (Fig. 3F, snATAC-seq tracks). Together, these observations indicate that *ERG* is a direct IKAROS target and that its repression in B-ALL reflects restoration of a developmental silencing program normally lost in *IKZF1*-mutated leukemias.

### IKAROS Proximal Proteome Reveals Association with ERG and Chromatin Regulators

Given that IKAROS limits ERG expression, we hypothesized that its tumor suppressor function may involve chromatin-regulatory complexes. To investigate these complexes, we performed TurboID proximity labeling in Ph B-ALL cells expressing N-terminal fusions of IK1 or IK6 (**S.** Fig. 4A-C). Mass spectrometry identified 196 high-confidence IKAROS-associated proteins enriched over empty vector controls (Table S8). IK1 and IK6 recovered highly similar sets of interacting proteins in TurboID experiments, as evidenced by comparable log -transformed mean intensities across identified proteins (Fig. 4A). Interactions included known IKAROS partners such as IKZF2, IKZF3, HDAC1, CHD4, and SIN3A, as well as a broader set of TFs (ERG, ZNF217, ZNF384), chromatin remodelers and co-regulators (DNMT1, SMARCA4, CTBP1, SMC1A), RNA-binding proteins (HNRNPA1, SND1, PRPF6), signaling mediators (PLCG2, TRIM25, YWHAB), and metabolic enzymes (FASN, ACLY, ACACA) (Fig. 4A). ERG was significantly enriched in both IK1 and IK6 TurboID datasets compared to empty vector controls (log fold change over controls = 1.48 and 1.83, respectively; p < 0.05; S Table 8).

**Figure 4.**
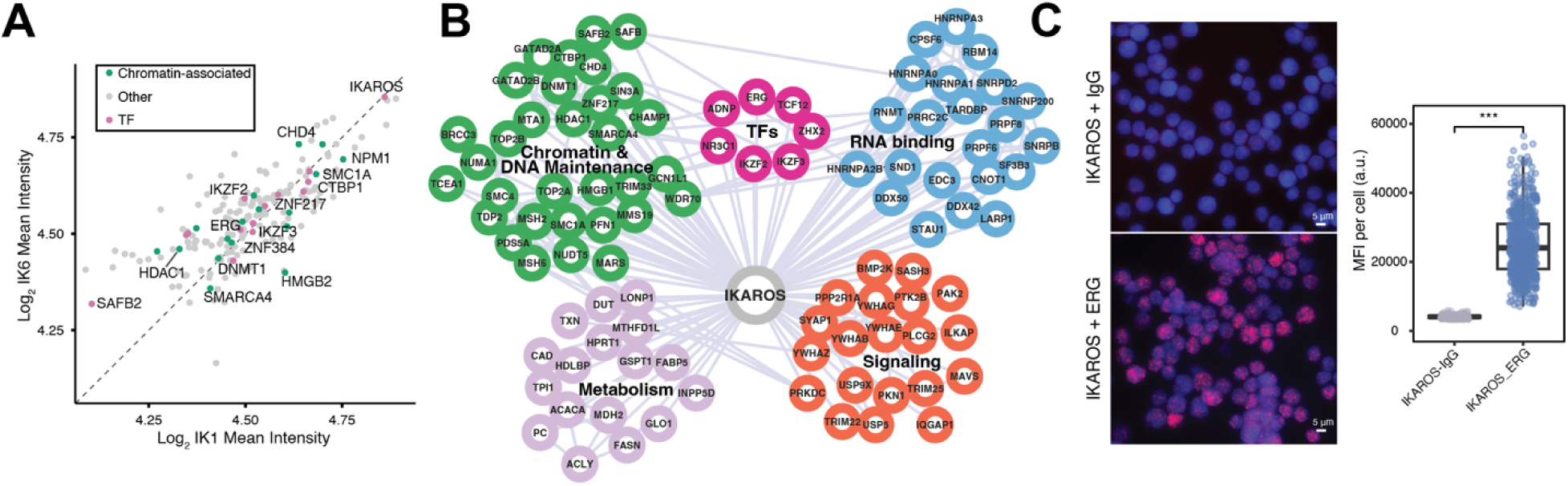
IKAROS proximity labeling reveals shared interactors between isoforms, including transcription factors, chromatin remodelers, and transcriptional co-regulators **A)** Comparison of log -transformed mean intensity of 196 high-confidence IKAROS-associated proteins identified by TurboID proximity labeling in Ph+ B-ALL cells expressing N-terminal TurboID fusions to I isoform 1 (IK1) or isoform 6 (IK6). Mean intensities averaged across biological replicates for each dataset. Proteins are colored according to curated functional annotations: transcription factors (pink), chromatin remodelers and co-regulators (green), and other proteins (gray). Selected known IKAROS interactions are labeled. **B)** Protein-protein interaction network of IKAROS-associated proteins derived from significantly enriched TurboID hits (Table S8). Nodes represent direct IKAROS proximal interactors grouped by functional categories: transcription factors (pink), chromatin/chromatin maintenance (green), RNA-binding proteins (blue), metabolic enzymes (purple), and signaling proteins (orange). Edges indicate high-confidence interactions among these proteins based on STRING database (score > 700). No second-degree interactors were included. **C)** Proximity ligation assay (PLA) confirms nuclear proximity association between IKAROS and ERG in MXP5 cells. Representative images (left) show PLA signal (red) and DAPI (blue) in cells stained with IKAROS and ERG antibodies (bottom) or IgG control (top). Quantification of mean fluorescence intensity (MFI) per nucleus (right) shows a significant increase in PLA signal in IKAROS–ERG samples compared to IgG controls (***p < 0.001, unpaired t-test, n = cells). Scale bar = 5 µm.

Co-immunoprecipitation using anti-HA-conjugated beads to capture HA-tagged TurboID–IKAROS fusions supported interactions with CHD4, ZNF217, and NRIP1 (Fig. S4D), whereas reciprocal immunoprecipitation of ERG did not recover IKAROS (data not shown). PLA further confirmed nuclear proximity between IKAROS and ERG in Ph B-ALL cells (Fig. 4C).

Functional annotation of the 196 IKAROS-associated proteins revealed significant enrichment for GO categories related to chromatin remodeling, nucleosome assembly, and DNA binding (adjusted p < 0.01; S Table 2). In addition to chromatin regulators, the network includes factors involved in RNA processing, metabolism, and signaling (Fig. 4B; Table S2). These findings support the interpretation that IKAROS interacts with a broad regulatory network, encompassing chromatin, signaling, metabolic, and RNA-processing functions. The enrichment of ERG among IKAROS-associated transcription factors, together with shared chromatin-modifying interactors, points to a mechanism by which IKAROS may influence ERG-dependent gene regulation.

Combined with the observed antagonism between IKAROS and ERG at shared chromatin targets (Fig. 2), these findings provide a mechanistic link between IKAROS-associated protein complexes and the transcriptional reprogramming induced by IK1 expression.

### IK1 induction remodels enhancer chromatin and alters ERG/IKAROS-associated regulatory states

We next examined how IK1 induction alters the chromatin landscape in MXP5 cells. We profiled histone modifications associated with enhancer activity (H3K27ac, H3K4me1), transcriptional elongation (H3K36me3), promoter activity (H3K4me3), and repression (H3K27me3, H3K9me3) under control and IK1 conditions. Differential enrichment was dominated by H3K27ac and H3K27me3, with >6,000 altered peaks in each mark, followed by ∼400 H3K9me3, and less than 150 H3K4me1, and H3K4me3 sites (Fig. 5A). The strongest TF overlap was observed for H3K27ac, where 27% of downregulated and 12% of upregulated regions intersected IKAROS binding, and ∼9–10% overlapped ERG (including ∼3% shared sites) (Fig. 5B). In contrast, overlaps with H3K27me3 and H3K9me3 peaks were ≤3%, and only a minority of the few H3K4me1 and H3K4me3 differential regions were associated with TF binding. These patterns indicate that IK1-dependent enhancer remodeling is most directly linked to ERG and IKAROS binding, whereas changes in Polycomb- and heterochromatin-associated marks potentially reflect indirect effects. Heatmaps and average profiles centered on differential peaks showed sharp, localized gains and losses of H3K27ac with IK1 induction, with IKAROS signal evident at both up- and downregulated regions and ERG signal highest in control cells at downregulated peaks (Fig. 5C). In contrast, H3K27me3 and H3K9me3 peaks displayed broader shifts in enrichment with little associated TF occupancy. We also used ChromHMM to segment combinatorial chromatin states and scored genome-wide transitions [37]. This revealed thousands of IK1-induced state shifts (Fig. S5), with the largest changes occurring at enhancer-and transcription-associated states. Together, these data indicate that IK1 induction primarily reshapes enhancer and promoter activity through direct IKAROS binding and displacement of ERG, while transitions involving Polycomb and heterochromatin states were less frequent and may reflect broader chromatin remodeling programs.

**Figure 5.**
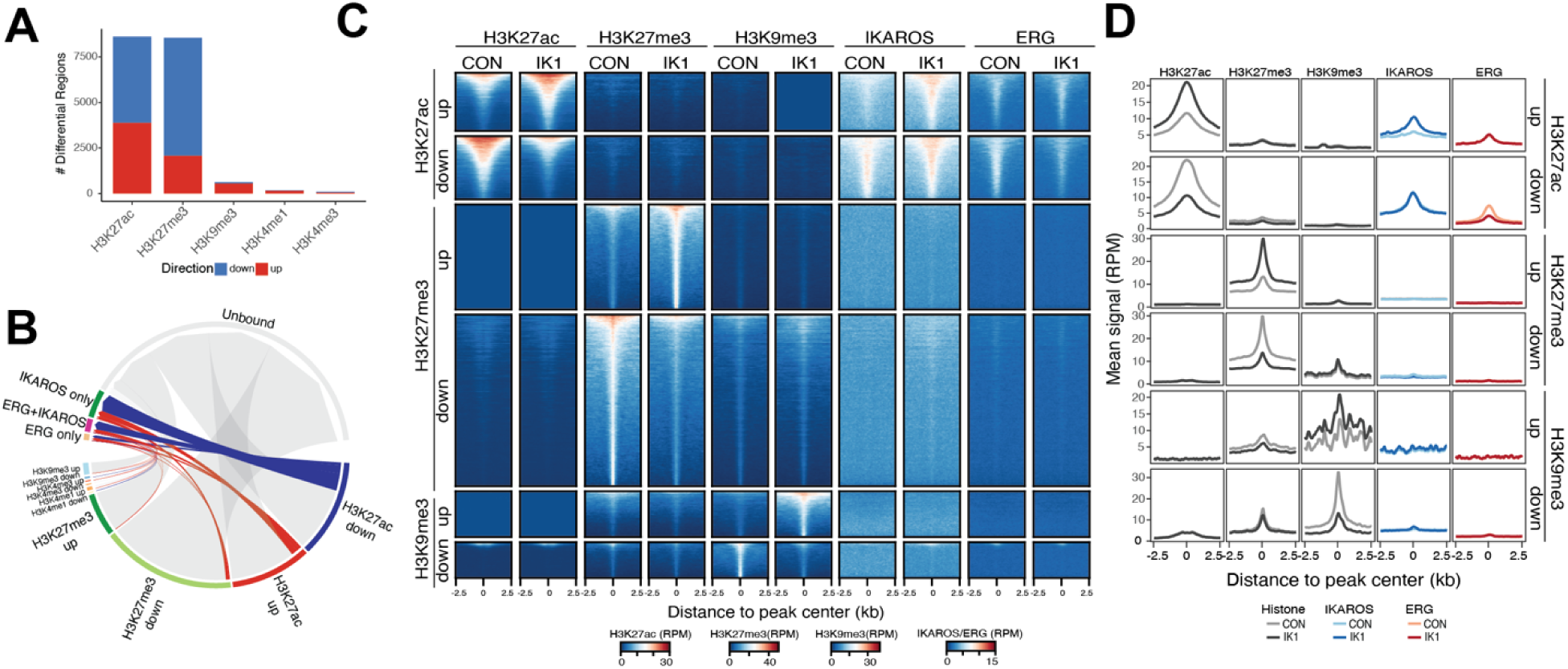
IK1 induction remodels enhancer chromatin and converges on ERG/IKAROS-bound sites. **A)** Differential analysis of histone modifications in MXP5 cells upon IK1 induction relative to control, showing the number of regions with significant changes (padj < 0.05, |log fold-change| > 1). Gains are shown in red and losses in blue. **B)** Relationship between IK1-induced changes in histone modification enrichment and TF binding. All regions with significant changes in H3K27ac, H3K27me3, H3K9me3, H3K4me1 and H3K4me3 (padj < 0.05, |log fold-change| > 1) are represented as arcs. Regions overlapping IKAROS-only, ERG-only, or shared IKAROS+ERG ChIP-seq peaks are shown in color, with gains in red and losses in blue. Differential regions that do not overlap IKAROS or ERG binding are shown in grey. **C)** ChIP-seq signal for H3K27ac, H3K27me3, and H3K9me3 at regions with significant changes (padj < 0.05, |log fold-change| > 1), displayed ±2.5 kb from the peak center and stratified by mark and direction of change (up or down). Heatmaps show normalized signal (RPM), with regions sorted by signal intensity. Corresponding IKAROS and ERG ChIP-seq profiles (RPM) are shown alongside the histone mark heatmaps. **D)** Aggregate signal plots for the same regions as in C, showing mean normalized ChIP-seq signal (RPM) for histone marks (gray), IKAROS (blue), and ERG (red) under control and IK1 conditions.

### IKAROS regulates enhancer-based spatial interactions

To determine whether IKAROS influences higher-order chromatin architecture through enhancer activity, we performed H3K27ac HiChIP in MXP5 cells under control and IK1-induced conditions with biological replicates. As expected for H3K27ac profiling, most detected loops corresponded to enhancer–promoter (E-P) or enhancer–enhancer (E-E) interactions (Fig. 6A).

**Figure 6.**
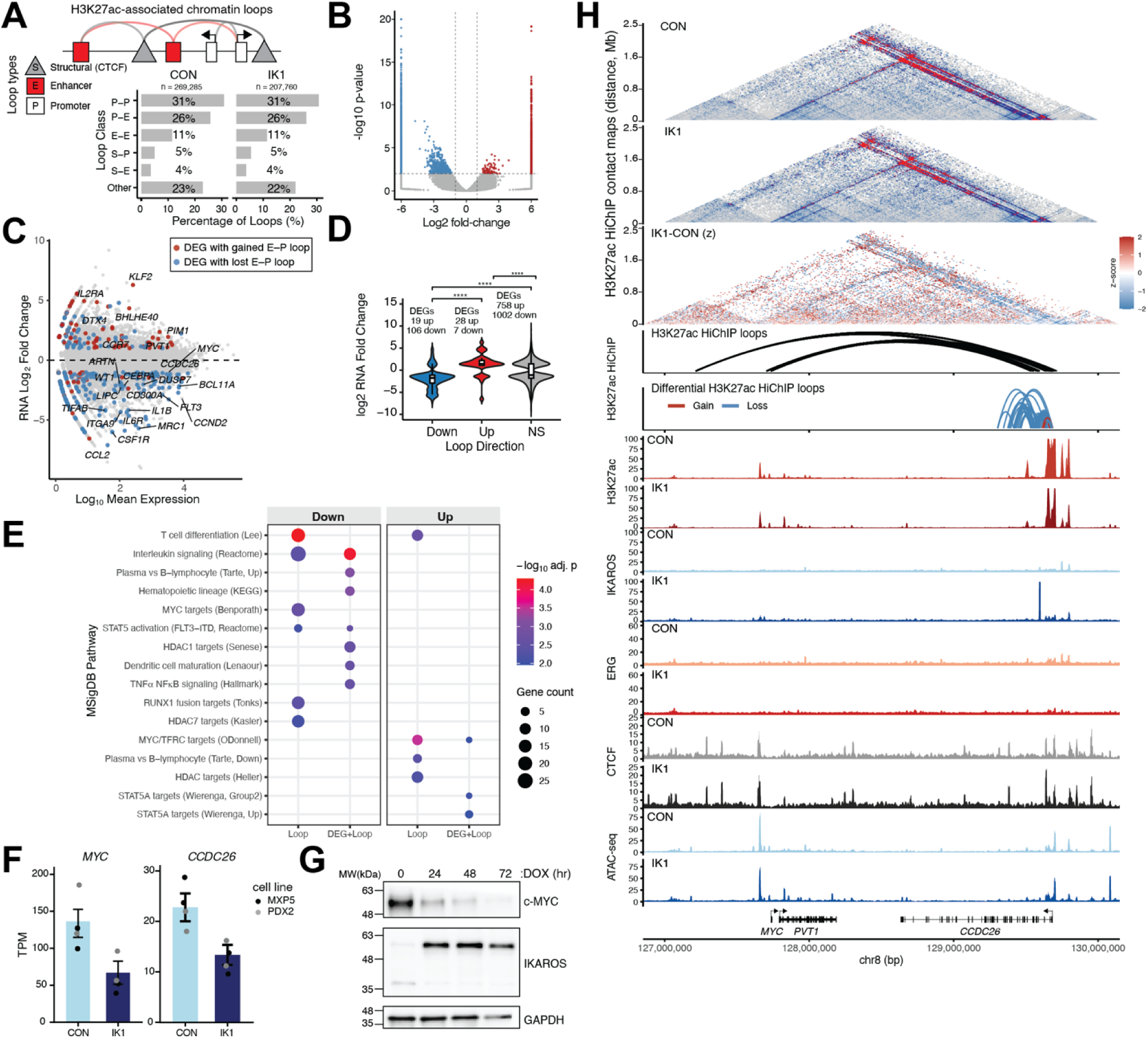
H3K27ac HiChIP identifies enhancer–promoter changes with IK1 induction. **A)** Distribution of loop types detected by H3K27ac HiChIP in control and IK1-induced MXP5 cells. P–P, promoter–promoter; E–P, enhancer–promoter; E–E, enhancer–enhancer; other. Table shows proportions of loop classes across conditions. **B)** Differential H3K27ac HiChIP loops (IK1 vs. CON). Loops with |log fold-change| > 1 and FDR < 0.05 were considered significant, with gains shown in red and losses in blue. **C)** RNA-seq differential expression plotted against mean expression, with genes linked to differential enhancer–promoter (E–P) loops marked. Blue, genes with gained loops; red, genes with lost loops. **D)** RNA-seq fold-change for genes linked to gained or lost E–P loops. Genes with loop gains show up-regulation, whereas those with loop losses show down-regulation. Statistical significance was assessed using a two-sided Wilcoxon rank-sum test with Benjamini–Hochberg correction, ****p < 1×10 . **E)** Pathway enrichment analysis of differentially expressed genes associated with gained (blue, left) or lost (red, right) enhancer–promoter loops. Enrichment was performed using MSigDB Hallmark (H) and Curated (C2) collections. Only pathways with adjusted p < 0.05 are shown. Dot size represents the number of genes contributing to each pathway; color encodes the adjusted p-value. **F)** RNA-seq expression (transcripts per million (TPM)) of *MYC* and *CCDC26* in MXP5 and PDX2 cells under CON (gray) and IK1 (black) conditions. Bars show mean ± SEM. **G)** Western blot of IKAROS and MYC protein levels in MXP5 cells after doxycycline (DOX)–induced IK1 expression (0–72 h). GAPDH was used as a loading control. **H)** HiChIP and ChIP-seq visualization at the MYC/PVT1/CCDC26 locus (chr8:127–130 Mb).

Differential loop analysis identified 2,122 significantly changed interactions (FDR < 0.05), including 1,510 lost and 612 gains following IK1 induction (Fig. 6B; Table S9). IK1-sensitive loops comprised E-P, E-E, and CTCF-associated contacts (Fig. S6C), with loop losses significantly longer than gains across E-P and structural–promoter (S-P) classes (Fig. S6D).

Integration with RNA-seq revealed concordant expression changes, where loop losses were associated with downregulation and loop gains with upregulation (Fig. 6C, D). Notably, *MYC* (log FC = –1.02, FDR = 3.3 × 10) and *CCND2* (log FC = –3.18, FDR = 1.0 × 10) exhibited transcriptional repression accompanied by loss of enhancer-linked loops, whereas *KLF2* (log FC = +6.30, FDR = 1.2 × 10 ³) and *BHLHE40* (log FC = +4.34, FDR = 5.4 × 10 ²) gained loops and were upregulated. Pathway analysis of loop-associated genes showed enrichment for MYC-target programs, hematopoietic differentiation, and cytokine signaling among downregulated genes with loop loss, whereas gained-loop genes were enriched for STAT5A activation and plasma-cell differentiation (Fig. 6E). At the *MYC* locus (chr8:127–130 Mb), which is subject to cell type–specific enhancer regulation across cancers [38–40], H3K27ac HiChIP revealed extensive E-P interactions in control cells linking *MYC* to *PVT1* and *CCDC26* (Fig. 6H). Following IK1 induction, contact frequency decreased across the region, with the most pronounced loss of interactions centered near the *CCDC26* promoter, which is also is also significantly downregulated (log₂FC = –1.03, FDR = 4.9 × 10⁻) (Fig. 6F, G). A comparable pattern was observed at *CCND2* (chr12:3.9–4.5 Mb), where H3K27ac HiChIP revealed loss of E-P interactions spanning the *CCND2-AS1* region following IK1 induction. These interaction losses coincided with reduced ERG occupancy and diminished IKAROS binding upon IK1 induction, consistent with observed IK1 repression of *CCND2* both at the RNA and protein levels (Fig. S7A-C). Locus-specific UMI-4C confirmed the HiChIP findings, showing significantly reduced contact frequencies from the *CCDC26* promoter and *CCND2* enhancer regions after IK1 induction (Fig. S7D). Together, these findings indicate that IK1 reshapes enhancer connectivity in a context-dependent manner, attenuating interactions at proliferative drivers such as *MYC*, *CCDC26*, and *CCND2*, while reinforcing enhancer networks linked to differentiation-associated programs.

Pooled H3K27ac HiChIP contact matrices from replicate experiments are shown at 10 kb resolution for CON (top) and IK1 (middle) conditions, followed by the differential HiChIP contact map (IK1 – CON). In the CON and IK1 panels, interaction strength is shown as z-score intensity (0–8). In the differential map, colors represent changes in interaction z-scores between conditions (red = gain, blue = loss) on a fixed z-score scale of −2 to 2. The y-axis represents genomic interaction distance (Mb), and the x-axis denotes genomic position on chromosome 8 (hg38). Black arcs show significant H3K27ac HiChIP loops detected in the pooled control H3K27ac HiChIP data (q ≤ 0.05) spanning 40–800 kb within the plotted region (chr8:127–130 Mb). The red and blue arcs represent significant H3K27ac HiChIP loop gains and losses (|log FC| ≥ 0.5, p < 0.05). Signal tracks show replicate-merged, RPKM-normalized ChIP-seq profiles for H3K27ac, IKAROS, ERG, and ATAC-seq, and MACS2 fold-enrichment tracks for CTCF, in CON and IK1 conditions.

### ERG is a functional dependency in *IKZF1*-deficient B-ALL

Given that *ERG* is a direct target of IKAROS and is strongly repressed upon IK1 re-expression, we next tested whether *ERG* contributes to leukemic maintenance in *IKZF1*-deficient B-ALL. We performed CRISPR interference (CRISPRi) growth competition assays using two independent sgRNAs targeting either the *ERG* TSS or a developmentally primed intronic enhancer within intron 3. This intronic region exhibits dynamic chromatin accessibility during B-cell development and is directly bound by IKAROS (Fig. 3F). Intracellular flow cytometry revealed robust reduction of ERG protein levels in mCherry (sgRNA-expressing) cells following TSS-targeting (also confirmed by western), and partial knockdown with intron 3-targeting guides in both PDX2 and TOM1 cells (Fig. 7A, B, S Fig. 8A). Quantification of ERG median fluorescence intensity (MFI) confirmed significant suppression across sgRNAs, with stronger knockdown observed for TSS-targeting constructs (Fig. 7C). Both targeting strategies impaired leukemia cell proliferation relative to non-targeting (NT) controls, though with distinct kinetics. In both cell lines, sgRNAs targeting the ERG TSS or intron 3 significantly reduced the frequency of mCherry cells, with effects first emerging between day 12 and 14 and persisting through day 31 (Fig. 7D). In TOM1 cells, ERG-g1 and ERG-g2 (TSS-targeting) and ERG-g5 (intron 3-targeting) significantly reduced mCherry cell frequency by day 14. ERG-g3 (intron 3-targeting) showed a more gradual depletion time course, reaching statistical significance by day 21. Growth suppression was considered significant at timepoints where both independent guides for a given region differed from NT-1 (FDR < 0.05). To assess the potential pharmacologic sensitivity of B-ALL cells to perturbation of ETS activity, a panel of lymphoid cell lines were incubated with the ETS family inhibitors YK-4-279 and its clinical derivative TK-216. Both compounds elicited dose-dependent reductions in viability across the tested panel, with half-maximal responses occurring around 1 μM (Fig. 7E). Finally, genome-scale CRISPR screening data from the Cancer Dependency Map revealed that *ERG* exhibits selective dependency in lymphoid malignancies, with B-ALL cell lines displaying the strongest and most consistent depletion phenotypes (Fig. 7F, S Fig. 8B). Together, these findings demonstrate that *ERG* contributes to leukemic maintenance in *IKZF1*-deficient B-ALL and highlight its potential as a lineage-restricted therapeutic vulnerability.

**Figure 7.**
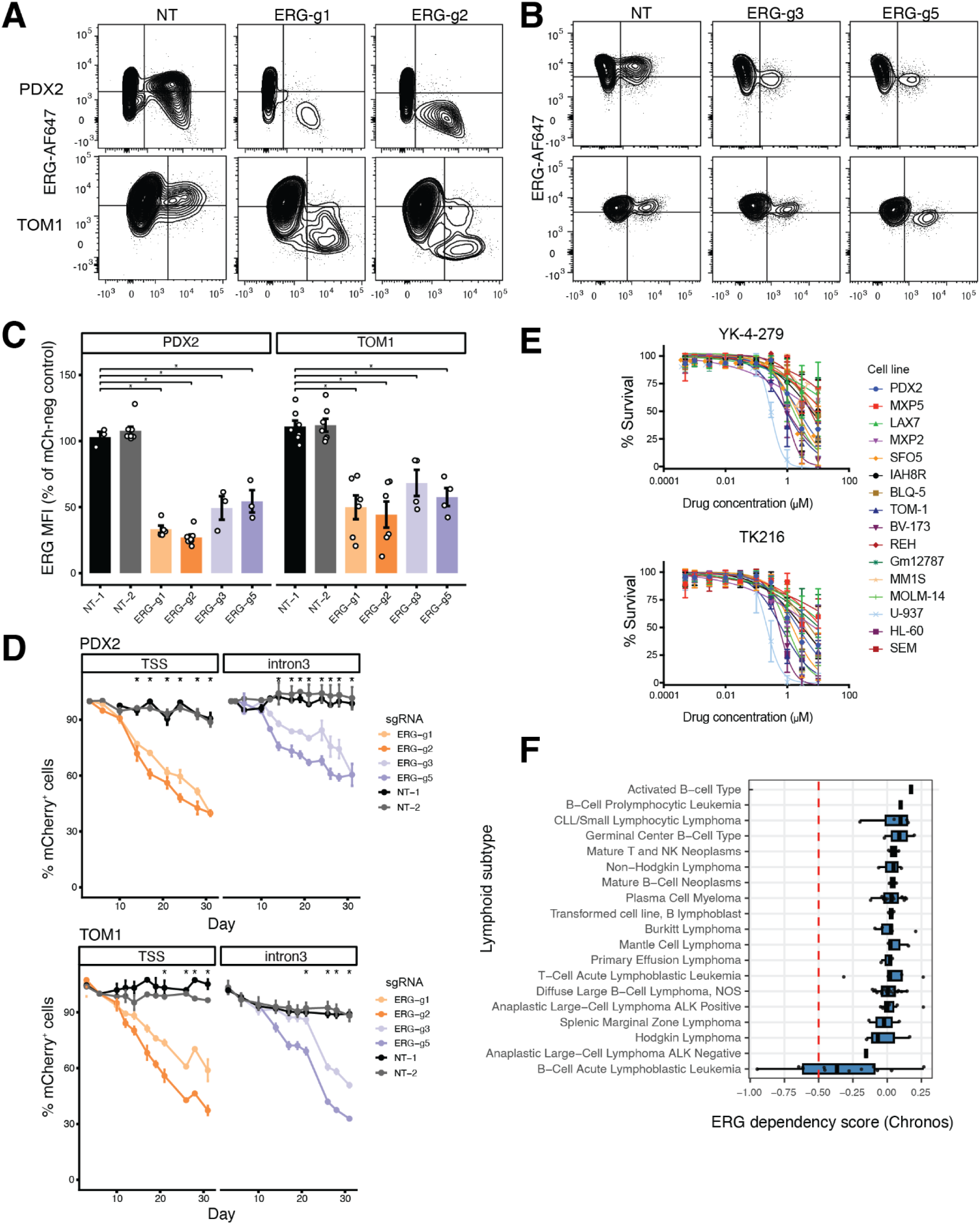
ERG Dependency of B-ALL for Cell Growth. **A and B)** Intracellular flow cytometry analysis of ERG protein levels following CRISPR interference (CRISPRi) targeting the *ERG* transcription start site (TSS) **A)** or a putative intronic enhancer **B)** in PDX2 and TOM1 cells. dCas9-KRAB^+^ (GFP^+^) cells were transduced with lentiviral sgRNA constructs co-expressing mCherry. Cells were fixed, permeabilized, and stained with ERG-AF647 antibody. Cells were pre-gated on singlets, live, GFP^+^ cells. Representative flow cytometry plots from day 6 post transduction display ERG protein levels (AF647) on the y-axis and mCherry signal on the x-axis. **C)** Quantification of ERG protein knockdown by intracellular flow cytometry. Median fluorescence intensity (MFI) of ERG-AF647 of the mCherry^+^ cells was normalized to the mCherry-negative population within each sample to control for staining variation. Bars represent mean ± SEM across biological replicates (PDX2: *n* = 3–8 per sgRNA; TOM1: *n* = 4–7 per sgRNA). Comparisons used unpaired two-sided t-tests vs non-targeting controls (NT) (per cell line), with Benjamini–Hochberg correction; * indicates FDR < 0.05. **D)** CRISPRi growth competition assays tracking the frequency of mCherry (sgRNA-expressing) cells over time in PDX2 and TOM1, normalized to day 3-4 post transduction. Two independent sgRNAs were used per target region (TSS or intron 3). mCherry frequency reflects the relative abundance of perturbed cells compared to non-targeted (sgRNA-negative) control cells within the same population. Curves show mean ± SEM from ≥ 2 independent replicates. Statistical significance was assessed at each timepoint using pairwise two-sided t-tests vs NT-1 (FDR-adjusted via Benjamini–Hochberg). * Indicates timepoints where both sgRNAs for TSS or intron 3 showed FDR < 0.05. **E)** Drug sensitivity of B-ALL and lymphoid cell lines to ETS inhibitors YK-4-279 and TK-216. Cells were treated with a range of concentrations for 48 hours, and viability was assessed using a resazurin-based assay. Absorbance values were normalized to untreated controls. Data represent mean ± SEM from ≥3 independent replicates. **F)** ERG dependency scores from the DepMap Chronos CRISPR dataset (22Q4). Lymphoid cancer cell lines are grouped by disease subtype. Each point represents a single cell line; boxplots indicate the median and interquartile range. The dashed red line marks the dependency threshold (Chronos score = –0.5).

## Discussion

Despite longstanding recognition of *IKZF1* as a tumor suppressor in B-ALL, the molecular mechanisms by which IKAROS enforces lineage restriction and constrains leukemic programs in humans have remained incompletely defined. Here, we used inducible expression of wild-type IK1 in *IKZF1*-deficient Ph B-ALL cells to dissect its early regulatory effects on chromatin and transcriptional networks. Multi-omic integration revealed that IKAROS re-expression remodels the B-ALL regulatory landscape through widespread changes in chromatin accessibility, enhancer activity, and transcription factor connectivity, notably repressing *ERG*, an ETS-family factor highly expressed in progenitor cells. This repression occurred alongside restoration of developmental regulatory programs characteristic of B-cell maturation, suggesting that *IKZF1* loss disrupts normal differentiation by failing to regulate progenitor-associated transcriptional circuits.

To test whether ERG contributes to leukemic cell growth and to identify regulatory elements controlling its expression, we targeted a developmentally active *cis*-regulatory element within *ERG* intron 3. This region displays open chromatin selectively during early B-cell differentiation in single-cell ATAC-seq analysis of normal development (Figure 2H and 3F) and becomes progressively inaccessible during development. Although prior studies have shown that *ERG* expression is developmentally regulated in hematopoiesis [41, 42], this intronic enhancer has not been previously characterized. We found that it is bound by IKAROS and undergoes chromatin changes in B-ALL cells closing upon IK1 induction, classifying it as a differentially accessible region. To test the functional relevance of this element, we used CRISPRi to target the intron 3 region, which significantly reduced *ERG* expression and impaired growth of different B-ALL cell lines (Figure 7), with effects comparable to CRISPRi targeting the *ERG* promoter region. These findings support a model in which IKAROS directly represses *ERG* through silencing of a developmentally primed *cis*-regulatory element in intron 3. The ERG dependency was further supported by pharmacologic inhibition. Treatment of B-ALL and lymphoid cell lines with ETS-family inhibitors YK-4-279 and TK-216 reduced cell viability at low micromolar concentrations, suggesting that ETS factor activity contributes to leukemic maintenance. YK-4-279 and TK-216 function by disrupting protein-protein interactions between ETS factors and essential cofactors, including the RNA helicases DHX9 (also known as RNA helicase A, RHA) and DDX5, thereby impairing transcriptional activation [43–45]. Although these compounds are not ERG-specific, they have demonstrated in vitro and in vivo antitumor activity in multiple lymphoma subtypes by disrupting ETS-helicase interactions [46]. In diffuse large B-cell lymphoma, this includes inhibition of SPIB and SPI1 function, leading to transcriptional repression of oncogenic programs and induction of apoptosis. Given the shared ETS dependence across B-lineage malignancies and the elevated ERG expression and ETS-associated gene regulatory networks observed in *IKZF1*-deficient B-ALL, our findings support the potential utility of these agents in selectively targeting ETS-related vulnerabilities in this context.

Restoring IKAROS function disrupts key enhancer–promoter interactions that support leukemic maintenance, particularly at loci co-occupied by ERG. ChIP-seq demonstrates that IKAROS and ERG co-occupy similar regulatory elements, including enhancers near *CCND2* and *CCDC26*, genes linked to proliferative and progenitor programs. *CCND2* encodes a G1 cyclin that is subject to activating mutations, translocations, and transcriptional upregulation in hematologic malignancies [47, 48], supporting its role as a potential effector of IKAROS-mediated enhancer repression. *CCDC26*, a long noncoding RNA located within the ∼2 Mb MYC locus and frequently amplified in leukemia [49], has been shown to regulate acute myeloid leukemia cell proliferation through KIT signaling and to influence erythroid differentiation via FOG-2–mediated transcriptional control of globin genes [50, 51]. Notably, *CCDC26* overlaps the BENC (blood enhancer cluster), a distal regulatory element that physically contacts the MYC promoter and is implicated in *MYC* activation via enhancer hijacking mechanisms in multiple hematologic malignancies [52]. At both loci, IKAROS binding is associated with changes in ERG occupancy, decreased H3K27ac, and diminished enhancer–promoter looping, consistent with enhancer decommissioning [38, 40, 53]. Together, these data highlight enhancer elements such as those at *CCND2* and *CCDC26* as direct targets of IKAROS-mediated repression, and potential mediators of leukemic maintenance in *IKZF1*-deficient settings.

Consistent with this model, IK1 induction increases IKAROS binding at these sites while ERG occupancy and H3K27ac levels decline, accompanied by weakened enhancer–promoter looping as shown by HiChIP and UMI-4C. In addition to chromatin co-occupancy, a novel and unexpected physical interaction between IKAROS and ERG was revealed by TurboID proximity labeling and validated by proximity ligation assays (Figure 4). This previously unreported interaction suggests that IKAROS may antagonize ERG function not only through competition for shared binding sites but also via proximal protein–protein interference. Complementing this, IKAROS was found to associate with chromatin remodeling complexes including NuRD, SIN3A–HDAC, and CHD4, that are known to modulate enhancer activity and chromatin structure [54–56]. The convergence of ERG interaction, chromatin remodeling, and enhancer architecture changes supports a model in which IKAROS displaces ERG from shared regulatory elements and recruits corepressors to regulate leukemogenic enhancer networks. In this context, IKAROS acts as a molecular scaffold, integrating sequence-specific binding, transcription factor antagonism, and chromatin modifier recruitment to silence oncogenic regulatory programs and enforce developmental gene expression fidelity.

Building on previous work demonstrating that IKAROS assembles lineage-specific 3D chromatin architecture in pre-B cells, including long-range enhancer to promoter interactions and the formation of superTADs that span TAD boundaries to organize regulatory units [27, 28], our study shows that re-expression of IKAROS in B-ALL cells partially reinstates this higher-order organization. Using orthogonal 3C methods, including HiChIP and UMI-4C, we find that IK1 reinstates enhancer–promoter loops at key differentiation-linked genes and regulates aberrant enhancer hubs that maintain oncogenic transcriptional programs. These architectural changes occur in the absence of CTCF redistribution and are accompanied by reduced chromatin accessibility and transcriptional downregulation at IKAROS-bound and ERG-bound regulatory loci. While IKAROS has been shown to organize chromatin topology during B-cell development [27], our study demonstrates its ability to reprogram diseased chromatin architecture in leukemia, reinforcing its role in both lineage specification and tumor suppression.

Beyond ERG antagonism, IKAROS exerts broader influence over lineage-restricted regulatory networks. Our gene regulatory network reconstruction revealed extensive redistribution of TF–TF and TF–target connectivity, shifting control from progenitor-associated ETS and E2F factors toward B-cell regulators such as PAX5, SPIB, and RUNX2. These IKAROS-driven network transitions recapitulate those observed during normal B-cell development, where ERG activity declines and IKZF1-associated regulons emerge. This developmental transition is consistent with in vivo degradation studies in mice demonstrating that Ikaros and Aiolos function primarily as transcriptional repressors during early B-cell commitment, acting in opposition to the activating roles of E2A, EBF1, and PAX5 [57].

Functionally, the suppression of ERG connects IKAROS-dependent chromatin remodeling to leukemic maintenance. ERG depletion or pharmacologic ETS inhibition lead to reduced proliferation and viability in B-ALL cells. These findings, together with clinical data showing that the adverse prognosis of *IKZF1* deletion is mitigated by co-occurring *ERG* loss, position ERG as a key effector of IKAROS deficiency and a potential therapeutic vulnerability in *IKZF1*-mutated leukemias. Although direct ERG targeting poses challenges due to its essential role in hematopoietic stem cells, combinatorial approaches can establish a leukemia-specific therapeutic window, and emerging approaches such as small-molecule degraders and disruption of ERG-stabilizing partners including USP9X may allow selective inhibition of aberrant ERG activity in leukemia. Consistent with a context-specific role in lymphoid malignancies, analysis of genome-scale CRISPR screening data from the Cancer Dependency Map (Fig. 7D; S Fig. 8) showed that *ERG* dependency is enriched in lymphoid lineages compared to other cancer types. Within lymphoid malignancies, B-ALL cell lines exhibited the strongest and most consistent depletion phenotypes, suggesting a selective requirement for *ERG* that may reflect dependence on progenitor-like transcriptional states.

Our study supports a central role for IKAROS as a regulator of enhancer topology, chromatin state, and transcriptional network hierarchy in Ph B-ALL. Through direct binding, recruitment of repressive complexes, and remodeling of 3D genome structure, IKAROS enforces developmental gene regulation and suppresses oncogenic enhancer programs driven by ERG and other progenitor-associated transcription factors. These findings clarify the mechanistic basis for *IKZF1* tumor-suppressor function and demonstrate that IKAROS restoration reestablishes lineage-specific chromatin architecture by coupling enhancer repression, transcriptional reprogramming, and genome organization into a unified model of leukemia restraint. Clinically, this model explains why the poor prognosis associated with *IKZF1* deletion is mitigated by co-occurring *ERG* loss, identifying ERG as a context-specific effector and therapeutic vulnerability. By linking IKAROS occupancy with enhancer architecture and 3D connectivity, IKAROS converts local chromatin engagement into higher-order regulatory outcomes that reinforce lineage fidelity and suppress leukemic potential. These findings expand the understanding of IKAROS from a DNA-binding repressor to a multifunctional architectural factor and suggest that restoring IKAROS-like chromatin states, or targeting their aberrant counterparts, may provide a tractable strategy for treatment of high-risk B-ALL.

## Methods

### Cell lines and culture conditions

Cells were obtained from Markus Müschen and DSMZ and listed in Table S10. Cells were maintained at 37°C in humidified incubator with 5% CO_2_ in minimum essential medium alpha (MEMα) GlutaMAX (Gibco, Thermo Fisher Scientific, Waltham, MA, USA), supplemented with 20% FBS (Gibco, Thermo Fisher Scientific, Waltham, MA, USA), tested negative for mycoplasma contamination and their identity was verified using STR profiling. IK1- and GFP-inducible cell lines were described previously[29, 30].

### Lentiviral production and transduction

HEK293T cells were transfected using MIRUS TransIT LT1 (Mirus Bio) with pCMV-dR8.91 and BaEVTR [58–60] packaging plasmids, along with the transfer plasmid. After 48 and 72 hours, medium was collected, filtered, concentrated 10x with Lenti-X concentrator (Takara Bio Inc), and either frozen for later use or mixed with 10^6^ cells in Retronectin-coated well (Takara Bio Inc.) in a 24-well plate and spun at 600-1000g for 30-45 minutes at 32°C.

### IK1 induction and flow cytometry

IK1 or GFP control expression was induced with 1 µg/mL doxycycline for 24 hours unless otherwise indicated. For assays requiring longer induction times, durations are specified in the corresponding subsections. When applicable, cells were analyzed by flow cytometry for GFP, IK1-linked reporters, or mCherry/marker expression depending on the assay. For intracellular flow analysis of ERG expression, cells were collected, incubated with Fc-block, stained with live/dead dye and washed with PBS between each step. Cells where then fixed for 10 minutes on 4C in the dark with 4% paraformaldehyde (Electron Microscopy Sciences). Cells were washed with PBS, and cell pellet was resuspended in Fixation/Permeabilization buffer for 20 min on ice and then washed with and stained in 1x Perm/Wash buffer according to manufacturer’s protocol (Invitrogen eBioscience Foxp3 / Transcription Factor Staining Buffer Set; Fisher Scientific).

Cells were resuspended in PBS and analyzed by flow cytometry. Antibody information is given in Table S10.

### Drug sensitivity assays

Cells were treated with varying doses of ETS family inhibitors YK-4-279 and TK216 for 48 hours. Cytotoxicity was measured using a resazurin assay (R&D Systems, AR002). Absorbance values were normalized to untreated samples, log transformed, fit to a nonlinear curve and plotted using GraphPad Prism.

### CRISPR interference

B-ALL cells were transduced with a dCas9-KRAB-P2A-GFP vector (Addgene #188771, gift from Marco Jost and Jonathan Weissman), packaged using pDR8.91 and BaEVTR helper plasmids [59, 60]. Lentivirus was produced in HEK293T cells as described above. GFP-positive cells were sorted and used for downstream experiments. sgRNAs targeting ERG, non-targeting controls, and a CD19 positive control were cloned into the pU6-sgRNA EF1α-Puro-T2A-mCherry vector (Addgene #217306, gift from Luke Gilbert). After transduction into dCas9-KRAB+ cells, mCherry-positive cells were quantified over time by flow cytometry. CD19 knockdown was used to validate repression efficiency. mCherry frequencies were normalized to 100% on day 3 or 4 post-transduction. Guide sequences are listed in Table S10. Statistical comparisons were made using unpaired two-tailed t-tests, with Benjamini–Hochberg correction.

### Western blotting

Cells were lysed in ice-cold RIPA buffer (50 mM Tris-HCl pH 7.4, 150 mM NaCl, 1% NP-40, 0.5% sodium deoxycholate, 0.1% SDS) supplemented with protease and phosphatase inhibitors (Roche Complete, Thermo Halt). Lysates were incubated on ice for 15 minutes with occasional vortexing, sonicated, and cleared by centrifugation at 14,000 × g for 10 minutes at 4 °C. Protein concentrations were determined using a BCA assay (Thermo Fisher). Equal amounts of protein were mixed with Laemmli buffer containing β-mercaptoethanol, boiled at 95 °C for 5 minutes, separated by SDS-PAGE, and transferred to PVDF membranes.

Membranes were blocked in 5% milk in TBS-T (0.1% Tween-20), incubated with primary antibodies overnight at 4 °C, then with HRP-conjugated secondary antibodies. Signals were detected using enhanced chemiluminescence and visualized by chemiluminescence imaging. Antibody information is provided in Table S10.

### Co-Immunoprecipitation

Cells expressing IK1 or IK6 in frame with miniTurbo-HA, or miniTurbo-HA alone, were induced with doxycycline for 48 hours, lysed in IP buffer (25 mM Tris-HCl pH 7.4, 150 mM NaCl, 1 mM EDTA, 1% NP-40, 5% glycerol, 2 mM MgCl) supplemented with BaseMuncher Endonuclease (abcam, ab270049) and protease inhibitors. Lysates were clarified by centrifugation at 16,000g for 10 min at 4°C. Supernatants were precleared with protein A/G magnetic beads (Pierce #88803) for 1 hour at 4°C. Supernatants were incubated with anti-HA magnetic beads (Pierce #88836) for 4 hours at 4°C with rotation. Beads were washed three times with wash buffer (TBS, 0.05% Tween), once with water and bound proteins were eluted with SDS sample buffer and analyzed by Western blotting. Antibody information is provided in Table S10.

### Proximity ligation assay

PLA was performed using the Duolink® In Situ Red Starter Kit Mouse/Rabbit (Sigma-Aldrich, #DUO92101) according to the manufacturer’s protocol. B-ALL cells were fixed and permeabilized in 100% methanol, then centrifuged onto microscope slides using a Cytospin™ centrifuge. Slides were incubated with primary antibodies listed in Table S10. Fluorescent signals were visualized using a Zeiss AxioImager epifluorescence microscope equipped with a Hamamatsu CCD camera. Images were captured at 40× magnification using Zen 2 software (Zeiss). Image analysis was performed in HALO software (Indica Labs) using the FISH module

(v3.1.3) or Highplex FL module (v3.2.1) to quantify PLA signal as dot counts per nucleus or total signal intensity. Data were exported and analyzed in R (v4.3.0). Nuclear segmentation was used to define individual cells, and per-cell MFI values were compared between conditions using a two-sided t-test.

### TurboID proximity labeling

A cassette containing miniTurbo, NLS and HA tag (amplified from Addgene #107172, kind gift from Alice Ting [61]) was cloned into TET-inducible pLVX vectors (Takara) containing IK1, IK6 or into an empty vector, followed by IRES and GFP. To create cell lines with stable and TET-inducible expression of miniTurbo constructs, plasmids were transduced into PDX2 cells expressing TET activator from pLVX-EF1a-Tet3G (Takara) Expression was assessed by Western blotting and biotinylation levels detected with Streptavidin-HRP (GenScript #M00091). Cells were induced with doxycycline for 24 hours and 50µM biotin was added for 30 minutes. Cells were lysed on ice in RIPA buffer (50 mM Tris-HCl (pH 7.5), 150 mM NaCl, 1% NP-40, 0.5% sodium deoxycholate, 0.1% SDS) supplemented with protease and phosphatase inhibitor cocktails (Roche). Lysates were clarified by centrifugation at 16,000 × g for 10 min at 4 °C. Supernatants were incubated with Dynabeads MyOne Streptavidin C1 magnetic beads (Thermo Fisher) at 4 °C for 3 h with gentle rotation. Beads were washed ×3, and bound proteins were eluted in SDS sample buffer, separated by 10% SDS-PAGE, and visualized by Coomassie staining. Gel lanes were excised, reduced with 10 mM DTT at 56 °C for 30 min, and alkylated with 55 mM iodoacetamide. Proteins were digested in-gel overnight at 37 °C with trypsin (Promega) in 50 mM ammonium bicarbonate. Peptides were extracted and resuspended in 0.1% formic acid and analyzed using a high-resolution Orbitrap mass spectrometer (Thermo Fisher) coupled to a nano-LC system. Raw MS data were processed using MaxQuant (v2.7.3.0) [62] with default settings (FDR 1%; LFQ intensities via MaxLFQ). Spectra were searched against the UniProt human reference proteome. Differential protein abundance was performed using the DEP2 R package (v 0.3.7.3) [63] and batch correction was applied using sva R package (v3.21) [64].

### Bulk RNA-seq library preparation and sequencing

Total RNA was isolated from 1–5 × 10 cells using the Direct zol RNA kit (Zymo Research). RNA integrity was confirmed with an Agilent Bioanalyzer, and concentration was determined using a Qubit fluorometer. Ribosomal RNA was depleted with the NEBNext rRNA Depletion Kit (Illumina), and libraries were prepared using the NEBNext Ultra II Directional RNA Library Prep Kit (NEB) with 1000 ng of input RNA. Libraries were sequenced on an Illumina HiSeq platform to generate single end 50 bp reads.

### Bulk RNA-seq analysis

Raw reads were assessed for quality using FastQC [65] and reads were aligned to the hg38 reference using STAR v2.7.11b [66] with GENCODE v36 annotation [67]. Gene level read counts were obtained using the --quantMode GeneCounts option in STAR. Differential expression analysis was performed using DESeq2 (v1.49.7) [68], with the design formula ∼ cell_line + condition to control for variation between MXP5 and PDX2 cell lines while testing for the effect of IK1 induction versus control. Genes with adjusted p-value < 0.05 (Benjamini–Hochberg correction) and absolute log fold change > 1 were considered differentially expressed genes (DEGs). Functional enrichment was assessed by Gene Set Enrichment Analysis (GSEA) using fgsea [69] against the MSigDB Hallmark and Canonical Pathways, pathways with FDR < 0.01 were considered significantly enriched [70, 71]. Genes were ranked by log fold change, and pathways with FDR < 0.01 were considered significant. Upstream regulator analysis was performed using Ingenuity Pathway Analysis (IPA) on significantly differentially expressed genes (adjusted p < 0.05) to identify potential transcriptional drivers. Regulators with predicted activation states and significant enrichment (p < 0.01) were reported. For analysis of public patient data, aligned RNA seq BAM files from Ph B ALL cases (EGAS00001007167 [34]) were downloaded. Gene level counts were computed using featureCounts [72]. *IKZF1* status (wild type vs. mutation) was defined as in Kim et al. [34], and differences in *ERG* expression between groups were evaluated using a Wilcoxon rank sum test.

### ATAC-seq library preparation and sequencing

ATAC-seq was performed using the opti-ATAC protocol [73] with 50,000 cells per condition, each with two independent biological replicates. Cells were washed with cold PBS and lysed on ice for 5 minutes in 50 μL of lysis buffer (10 mM PIPES pH 6.8, 100 mM NaCl, 300 mM sucrose, 3 mM MgCl , 0.1% Triton X-100, protease inhibitors). Nuclei were pelleted at 2500 rpm for 5 minutes and resuspended in 20 μL of transposition mix (10 μL TD buffer, 0.8 μL Tn5 transposase (Illumina, 9.2 μL water). Transposition was performed at 37°C for 30 minutes. Tagmented DNA was purified using the MinElute PCR Purification Kit (Qiagen), and libraries amplified using NEBNext High-Fidelity PCR Master Mix with barcoded Nextera primers. Libraries were double size-selected with AMPure XP beads (Beckman Coulter) to retain fragments between 150–1000 bp, quality-checked on an Agilent Bioanalyzer, and sequenced as paired-end 150 bp reads on an Illumina NovaSeq platform.

### ATAC-seq analysis

Raw reads were aligned to the hg38 reference genome using the BWA-mem aligner [74]. PCR duplicates were removed and ENCODE blacklisted regions were excluded. Peaks were called using MACS2 v2.2.9 [75], and a consensus peak set was created from merged replicates. Read fragments were counted per peak to generate a peak count matrix. Peak counts were normalized and variance-stabilized using DESeq2 [68]. PCA was performed on the 500 most variable peaks. Differentially accessible regions (DARs) were identified using DESeq2, with significance thresholds of adjusted p-value < 0.05 and |log fold change| > 1. TF footprinting was performed using the TOBIAS pipeline [31]. BAM files were corrected using ATACorrect and footprint scores were computed using ScoreBigwig. Differential TF binding scores between IK1 and control conditions were computed using BINDetect, with motifs showing absolute binding score difference ≥ 0.2 and p-value < 0.05 considered significant. Motif logos were generated from predicted bound regions using universalmotif [76].

### ChIP-seq library preparation and sequencing

ChIP-seq was performed following a modified protocol from the Farnham laboratory [77]. Cells were fixed in 1% formaldehyde for 10 minutes at room temperature, quenched with 125 mM glycine, washed twice with cold PBS, pelleted, and snap-frozen. Nuclei were lysed in buffer (50 mM Tris-Cl pH 8.1, 10 mM EDTA pH 8.0, 1% SDS, protease inhibitors) for 30 minutes, followed by one freeze–thaw cycle. Chromatin was sheared to an average fragment size of 200–500 bp using a Covaris S220 sonicator. For each immunoprecipitation, 50 µg of chromatin was incubated overnight with a primary antibody (**S** **Table 10**) and captured with 50 µL of protein A/G magnetic beads (Pierce #88803). After sequential washes and elution, cross-links were reversed, and DNA was purified using the PureLink PCR Purification Kit (Invitrogen #K310001). Input chromatin (500 ng) was processed in parallel. Libraries were prepared using the NEBNext Ultra II DNA Library Prep Kit (NEB #E7645) and sequenced as single end 50 bp reads on an Illumina HiSeq 1500 platform.

### ChIP-seq analysis

Raw reads were assessed using FastQC [65] and adapters were trimmed prior to alignment. Reads were aligned to the human genome (hg38) using STAR v2.7.11b [66] in end to end mode with intron spanning alignments disabled. Duplicate and low quality reads were removed using samtools [78]. Peak calling was performed using MACS2 v2.2.9 [75] with matched input controls and a q value threshold of 0.01. Fold enrichment signal tracks were generated from the aligned BAM files using UCSC bedGraphToBigWig utilities [79]. Replicate reproducibility was evaluated using the Irreproducible Discovery Rate (IDR) framework [80] and high confidence peaks were defined at IDR < 0.01 using IDR2D R package [81]. QC metrics including mapped read counts, peak counts, and fraction of reads in peaks (FRiP) were computed using ssvQC / seqsetvis v1.28 [82]. Peak signal distribution and read coverage were quantified from read per million (RPM) normalized BAM files across peak regions using deepTools v3.5.0 [83] and seqsetvis. Peak annotations (promoter, intragenic, intergenic) were generated using ChIPseeker v3.21 [84] with GENCODE v36 gene models [67]. Overlaps between ChIP-seq peak sets across control and IK1 conditions were computed using GenomicRanges v1.60.0 [85]. Consensus peak sets for each factor and condition were defined from IDR filtered peaks and used for downstream analyses. Genomic annotation of peaks was performed using ChIPseeker with TxDb.Hsapiens.UCSC.hg38.knownGene and org.Hs.eg.db annotation packages. Peaks were classified as promoter, intragenic, or distal intergenic based on overlap with transcription start sites (±3 kb), gene bodies, or intergenic regions. Motif enrichment analysis was performed using the monaLisa [86] Bioconductor package (v4.5). Each peak set was resized to a 500 bp window centered on peak summits and corresponding genomic sequences were extracted using the BSgenome.Hsapiens. UCSC.hg38 reference genome.

Vertebrate position weight matrices were retrieved from the JASPAR2020 database [87] via TFBSTools (v3.22) [88] and converted into PWMs for scanning. Motif enrichment was quantified using calcBinnedMotifEnrR() from monaLisa, using bin-level composition-corrected enrichment modeling, with default parameters and SerialParam(). Enrichment results were grouped by motif families based on motif names using custom regular expressions. For each family, the motif with the maximum enrichment score in at least one group was retained. Log enrichment scores were visualized using ComplexHeatmap (v3.21) [89]. Motif clustering was performed using Pearson similarity of position frequency matrices and displayed as dendrograms with dendrogram-informed heatmaps alongside representative motifs. Positional motif analysis was performed using curated PWMs for ERG and IKZF1 from MotifDb (v1.51.0) [90], selecting top motifs from HOCOMOCO [91] and JASPAR [87]. Peaks were resized to 400 bp windows around the summit and scanned using getPwmMatches() with a 70% match threshold and best central match retained. For each match, the relative distance to the peak center was calculated and binned in 20 bp intervals. Motif densities were normalized by peak set and visualized using ggplot2 (v4.0.0) [92] with smoothed line plots.

### CUT&RUN

The CUTANA CUT&RUN kit (EpiCypher #14-1048) was used according to manufacturer’s instructions, using 500,000 cells per reaction. Briefly, Concanavalin A beads were used to immobilize cells and .5 μg of individual antibodies against H3K4me3 (Epicypher #13-0041), H3K27me3 (Active Motif, #39159), and H3K9me3(Active Motif, #39161) were incubated overnight at 4°C with light rocking using 1 μL control IgG (Epicypher #13-0042) antibodies. Antibody chromatin complexes were treated with pAG-MNase for 2 hours at 4°C and DNA fragments were purified. DNA concentration was measured with a Qubit fluorometer and 5-10ng of purified DNA was input into sequencing library preparation with NEBNext UltraII kit (NEB # E7645). Library size was assessed with an Agilent Bioanalyzer and sequenced paired-end 150bp on an Illumina platform.

### CUT&RUN analysis

CUT&RUNTools analysis pipeline was used for processing [93]. Briefly, reads were trimmed and aligned to the hg38 genome. Narrow peaks were called from deduplicated reads with MACS2 and ssvQC was used to assess signal enrichment over IgG control. seqsetvis was used for visualization of genomic signal [82].

### Chromatin state modeling (ChromHMM)

Chromatin state segmentation was performed using ChromHMM (v1.27) [37] to define combinatorial regulatory states across the genome. CUT&RUN and ChIP-seq datasets were generated for six histone modifications (H3K27ac, H3K4me1, H3K4me3, H3K27me3, H3K9me3, H3K36me3) and CTCF in MXP5 cells under control and IK1-induced conditions. BAM files for each biological replicate were binarized separately at 200 bp resolution using ChromHMM BinarizeBam function, with matched IgG or input control files. All binarized replicates were included to learn a 15-state model using the LearnModel function. Emission probabilities were used to annotate each state based on canonical histone mark and CTCF signatures. For state annotation, genome-wide segment enrichments were calculated against the ENCODE Roadmap 18-state reference model using hematopoietic and B cell–derived epigenomes [94]. For each ChromHMM state, overlap widths with each ENCODE state were computed, normalized to genome coverage, and used to derive fold enrichment and maximum fractional overlap. MXP5 states were then assigned to functional categories based on both emission signatures and the most enriched ENCODE references. Annotated state groups included active promoters (S1–S3), enhancers (S4–S7), transcribed gene bodies (S8), heterochromatin (S9–S10), CTCF-associated boundaries (S11), Polycomb-repressed regions (S12), bivalent chromatin (S13), and quiescent/low-signal regions (S14–S15).

### Dynamic chromatin domain analysis (SCIDDO)

Dynamic chromatin domains (DCDs) were identified using SCIDDO (v1.0) [95]. BED-formatted ChromHMM segmentations for control and IK1-induced samples and the corresponding model file were used as input. SCIDDO was run with default settings, and segments with positive differential scores were merged and retained as dynamic chromatin domains. Domains were annotated by their ChromHMM state in control and IK1 conditions.

### Gene regulatory network analysis

Gene regulatory networks were inferred using the Paired Expression and Chromatin Accessibility (PECA) statistical framework [32]. Transcript abundance was quantified using Salmon [96]. Paired TPM matrices and ATAC-seq BAM files were used as input to PECA with the hg38 genome as reference. For each condition (IK1-induced and control, or *IKZF1*-mutant vs *IKZF1*-wt in Supplemental Figure 1), a network of TF–TG regulatory interactions were inferred. Interactions were considered condition-specific if present in one condition but not the other. To assess TF activity within conditions, networks were filtered to retain interactions with PECA scores > 0.5, and normalized degree centrality was calculated using the igraph R package [97].

The mean normalized degree for each TF was computed across replicates to estimate relative regulatory influence. To compare networks across conditions, the multi-sample differential PECA function was used. The resulting GRNs were filtered to retain high-confidence edges with TF–TG correlation > 0.3. Final networks were visualized using igraph following additional degree-based filtering.

### Single Cell multiome analysis

Paired scRNA-seq and scATAC-seq data from healthy donor bone marrow were obtained from GEO (GSE194122). Raw FASTQ files were processed with Cell Ranger ARC (10x Genomics, GRCh38 reference 2024-A) to generate gene-barcode matrices and ATAC fragment files. Cells with outlier RNA or ATAC metrics were removed during preprocessing using Scanpy [98]. For RNA, cells with fewer than 500 detected genes, fewer than 1,000 UMIs, or greater than 15% mitochondrial RNA content were excluded. For ATAC, cells with fewer than 1,000 unique fragments, fewer than 20% fragments-in-peaks, or a TSS enrichment score below 6 were excluded. Putative doublets were identified as outliers in joint RNA/ATAC distributions and removed. Thresholds were further adjusted per donor based on inflection points in the quality metric distributions. For the RNA modality, expression matrices were library-size normalized using log1p transformation, and highly variable genes were selected. Principal component analysis was performed on the top 50 components, and the first 30 PCs were used to construct a k-nearest neighbor graph. Uniform manifold approximation and projection (UMAP) embeddings were generated, and clustering was performed using the Leiden algorithm at a resolution parameter of 0.5. Hematopoietic progenitor and B-lineage subsets (HSC, CLP, Pro-B, Pre-B, Immature B) were annotated using canonical marker expression, including *CD34* and *SPINK2* for HSC, *TCF3* and *DNTT* for *CLP*, *VPREB1*, *IGLL1*, and *PAX5* for Pro-/Pre-B, and *MS4A1* and *IGHM* for immature B. To visualize developmental trajectories, the medoid coordinates of each stage were connected by a cubic spline in the biologically ordered sequence from HSC through immature B. For each TF group, per-cell module scores were computed as the average expression of the TF set minus the mean of a matched control gene set (Scanpy tl.score_genes, ctrl_size = 50, n_bins = 25; Seurat AddModuleScore with default settings). Scores are relative enrichments and were summarized by discrete developmental stages. For regulatory network inference, eRegulons and their activities were inferred with SCENIC+ [33], integrating motif accessibility and target-gene co-expression. Activity was summarized as +/+ (activated), −/− (repressed), or −/+ (discordant). Developmental dynamics were visualized on pseudotime computed from the RNA kNN graph. All analyses were performed in Python, using Scanpy for preprocessing/UMAP and SCENIC+ for regulon inference/visualization.

### HiChIP library preparation and sequencing

HiChIP was performed using the HiC+ kit (Arima A101020), according to the manufacturer’s protocols, with slight modifications. Cells were crosslinked with 1% formaldehyde for 10 minutes at room temperature, quenched with 125mM glycine. HiC was performed using the Arima protocol with 15 μg of chromatin per sample. Chromatin was then sheared to average size of 500 bp using Covaris S220 sonicator, followed by incubation with 4 μl of H3K27Ac antibody (Active Motif, 39135) overnight and chromatin immunoprecipitation with protein A/G magnetic beads (Pierce 88803), followed by washes according to the Arima protocol. Capture of biotinylated fragments and library building was performed using Swift Biosciences Accel-NGS 2S Plus DNA Library Kit (Cat # 21024) according to manufacturer’s instructions. Libraries were sequenced as paired end 150bp on Singular Genomics G4 sequencer.

### HiChIP analysis

HiChIP analysis was performed using the HiC-Pro pipeline [99] to generate valid interaction pairs from Bowtie2-aligned reads (hg38). Only uniquely mapped, non-duplicate reads from valid ligation products were retained. Downstream interaction analysis was performed using FitHiChIP (v11.0) [100] with peak-to-all interactions (IntType=3) at 10 kb resolution. Loops were called within a distance range of 20 kb to 4 Mb, and significant interactions were identified at FDR < 0.01 using coverage bias regression (BiasType=1) and merging of nearby interactions (MergeInt=1). MXP5 H3K27ac peaks were used as anchors. FitHiChIP was run in both replicate and pooled modes across conditions.

### UMI-4C library preparation and sequencing

UMI-4C libraries were generated from 10 million MXP5 cells collected 24 hours after doxycycline induction, following a modified protocol [101]. Briefly, cells were crosslinked with 1% formaldehyde for 10 minutes, quenched with 0.125 M glycine, washed with ice-cold PBS, and pelleted. Nuclei were isolated using freshly prepared lysis buffer (50 mM Tris-HCl pH 7.5, 150 mM NaCl, 5 mM EDTA, 0.5% NP-40, 1% Triton X-100, and protease inhibitors) and homogenized using a glass dounce on ice. Nuclei were then resuspended in NEB DpnII buffer supplemented with SDS and incubated at 37 °C with shaking. Triton X-100 was added to sequester SDS, and chromatin was digested with high-concentration DpnII in three rounds (2 h, overnight, and 2 h at 37 °C). Digestion efficiency was assessed by gel electrophoresis following Proteinase K digestion and phenol-chloroform extraction. Enzyme was inactivated at 65 °C, and proximity ligation was performed overnight at 16 °C using T4 DNA Ligase. After reversing crosslinks, 3C DNA was purified by phenol-chloroform extraction and ethanol precipitation. Purified 3C DNA was sonicated to ∼400-600 bp using a Covaris S220. Fragment size was confirmed by Bioanalyzer (Agilent). DNA was end-repaired (NEBNext E6050L), cleaned with 2.2× Ampure XP beads (, A-tailed using Klenow (NEB M0212M), and dephosphorylated with calf intestinal alkaline phosphatase (NEB M0290S). Forked, Illumina-compatible indexed adapters were custom-annealed from equimolar top and bottom oligos in annealing buffer and ligated using Quick Ligase (NEB M2200) at 25 °C. Unligated strands were removed by denaturation at 95 °C for 2 minutes, followed by 1× Ampure XP cleanup.

Two rounds of nested PCR were used to enrich for interactions with specific viewpoints. The first PCR used an upstream (US) bait-specific primer with an Illumina enrichment primer; the second used a downstream (DS) bait-specific primer carrying a P5 overhang. Libraries were purified with Ampure XP beads (1× and 0.7×), quantified with Qubit (Thermofisher), and fragment size confirmed by Bioanalyzer. Libraries were generated from at least two independent 3C templates (with distinct barcodes), pooled from 5–10 PCR reactions per template to increase complexity and allow technical replication. Adapter and primer sequences are listed in Table S10.

### UMI-4C data analysis

Chromatin interactions and differential contact analysis between IK1-induced and control cells were performed using the UMI4Cats R package (v1.20.0) [102]. Raw sequencing reads were first filtered by matching the sequence between the second-round PCR primer and the DpnII restriction site to retain only those originating from the designated viewpoint. The sequence downstream of the restriction site was mapped to DpnII-digested hg38 to identify interacting fragments. Reads were collapsed using unique molecular identifiers (UMIs) derived from random sonication ends. Only fragments within 3 kb to 2 Mb of the viewpoint were retained. UMI counts were variance-stabilized and smoothed, and chromatin interactions were called based on Z-scores and false discovery rate calculations. Differential interactions between conditions were identified using Fisher’s exact test, excluding regions with fewer than 10 total UMIs.

### DepMap analysis

The CRISPR (DepMap Public 23Q2+ Chronos Score) dataset was downloaded from the DepMap portal (https://depmap.org/portal/) for 1096 cell lines [103, 104]. Lineages containing less than 5 cell lines were removed from the analysis.

## Data availability

The total RNA-seq, ATAC-seq, ChIP-seq, CUT&RUN and HiChIP data reported in the present study are available at the Gene Expression Omnibus (GEO) repository under accession no. GSE310976.

## Acknowledgements

We thank Luke Gilbert, Marco Jost, Jonathan Weissman, and Markus Müschen and for kind gifts of plasmids and cells, respectively, and members of the Frietze and Schjerven lab for technical support. We acknowledge the Parnassus Flow Cytometry CoLab at UCSF (RRID:SCR_018206) supported in part by Grant NIH P30 DK063720 and by the NIH S10 Instrumentation Grant S10 1S10OD026940 01, the Harry Hood Bassett Flow Cytometry and Small Particles Detection facility (RRID:SCR_022147) and the Vermont Integrative Genomics Resource Facility (RRID: SCR_021775) at UVM, supported by an Institutional Development Award (IDeA) from the National Institute of General Medical Sciences of the National Institutes of Health under grant number P20GM103449, and U54 GM115516 for the Northern New England Clinical and Translational Research Network. This work was supported by grants from National Institutes of Health, R21CA209229 and R01CA230618 (S.F. and H.S.). Its contents are solely the responsibility of the authors and do not necessarily represent the official views of NIH.

