## Supplemental Figures for "IKAROS Gene Regulatory Network Reveal ERG as a Vulnerability in B-cell Acute Lymphoblastic Leukemia"

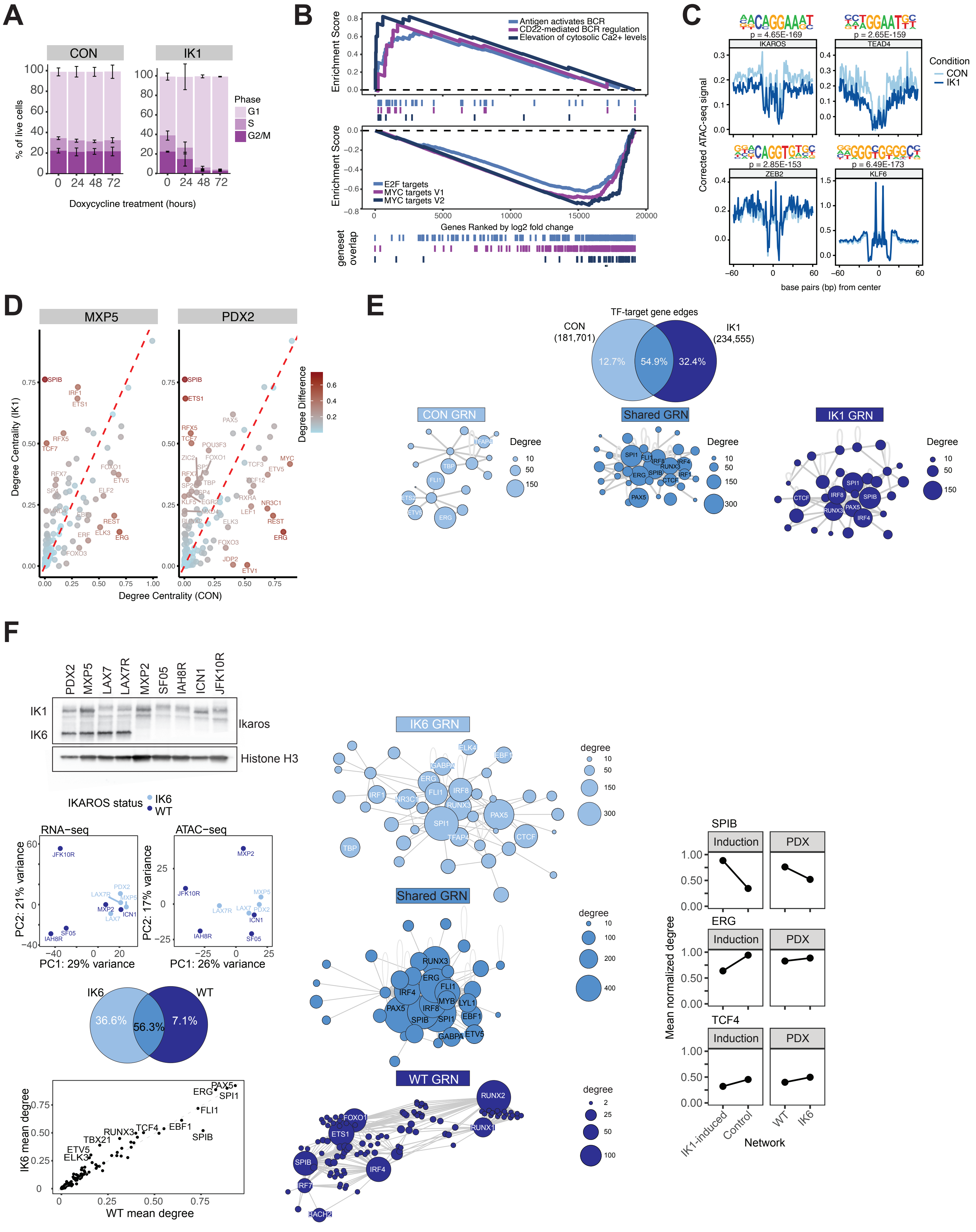
A
CON
IK1
% of live cells
Phase
G1
S
G2/M
Doxycycline treatment (hours)
B
Enrichment Score
Antigen activates BCR
CD22-mediated BCR regulation
Elevation of cytosolic Ca2+ levels
E2F targets
MYC targets V1
MYC targets V2
Genes Ranked by log2 fold change
geneset overlap
C
p = 4.65E-169
IKAROS
p = 2.65E-159
TEAD4
p = 2.85E-153
ZEB2
p = 6.49E-173
KLF6
Corrected ATAC-seq signal
Condition
CON
IK1
base pairs (bp) from center
D
MXP5
PDX2
Degree Centrality (IK1)
Degree Centrality (CON)
Degree Difference
E
TF-target gene edges
CON (181,701)
IK1 (234,555)
12.7%
54.9%
32.4%
CON GRN
Shared GRN
IK1 GRN
Degree
F
PDX2
MXP5
LAX7
LAX7R
MXP2
SF05
IAH8R
ICN1
JFK10R
IK1
IK6
Ikaros
Histone H3
IKAROS status
IK6
WT
RNA-seq
ATAC-seq
PC2: 21% variance
PC1: 29% variance
PC2: 17% variance
PC1: 26% variance
IK6
WT
36.6%
56.3%
7.1%
IK6 mean degree
WT mean degree
IK6 GRN
Shared GRN
WT GRN
degree
SPIB
ERG
TCF4
Induction
PDX
Mean normalized degree
IK1-induced
Control
WT
IK6
Network

**Figure S1. IKAROS re-expression alters chromatin accessibility and rewires gene regulatory networks in B-ALL cells.**

**A)** Cell cycle analysis by PI staining of MXP5 cells following doxycycline-induced IK1 expression. Proportion of cells in each cell cycle phase (G1, S, G2/M) was quantified at indicated timepoints. Data is replotted from previously published data [29].

**B)** Gene set enrichment analysis (GSEA) of RNA-seq data following 24-hour IK1 induction. Top: Enrichment of immune signaling pathways including B-cell receptor and calcium signaling. Bottom: Enrichment plots for MYC and E2F targets, which are downregulated with IK1 expression.

**C)** Representative TOBIAS transcription factor footprinting profiles for selected motifs. Aggregated ATAC-seq signal around motif centers (±60 bp) in control (light blue) and IK1-induced (dark blue) samples reflect differential TF occupancy. Motif logos and statistical significance are shown.

**D)** Condition-specific GRN rewiring across the two individual cell lines. Degree centrality of transcription factors in control versus IK1 networks in MXP5 (left) and PDX2 (right). Each point represents a TF, with position indicating GRN degree (i.e., connectivity to target genes) under control (x-axis) and IK1 (y-axis). TFs above the diagonal (red dashed line) show increased connectivity after IK1 induction, while those below exhibited higher target gene connectivity in CON. Color scale reflects magnitude of connectivity change.

**E)** Comparative analysis of gene regulatory networks in *IKZF1* wild-type (WT) and *IKZF1*-mutated ($IK6^{+}$) B-ALL cell lines. Top left: Western blot analysis of Ikaros protein expression status with full-length (IK1) and dominant negative (DN) IK6 indicated. Middle: PCA of RNA-seq (left) and ATAC-seq data (right) separates IK6$^+$ and WT lines. Bottom: Scatterplot of mean TF degree in WT versus IK6$^+$ GRNs shows widespread network differences.

**F)** *IKZF1* status is associated with distinct GRN configurations in a panel of B-ALL cell lines. Top left: Immunoblot of IKAROS protein in B-ALL cell lines showing full-length IK1 and the IK6 isoform. Histone H3 was used as a loading control. Middle left: Principal component analysis of RNA-seq (left) and ATAC-seq (right) data across *IKZF1*-mutant (IK6; light blue) and wild-type (WT; dark blue) cell lines. Bottom left: Scatterplot comparing mean TF degree (connectivity) between WT and IK6 networks. Center: GRN layouts showing TF-target edges specific to IK6, shared between IK6 and WT, or specific to WT. Node size reflects TF degree. Right: Normalized TF connectivity (mean degree) for SPIB, ERG, and TCF4 across IK1 induction (MXP5, PDX2) and B-ALL cell lines stratified by IKZF1 status.

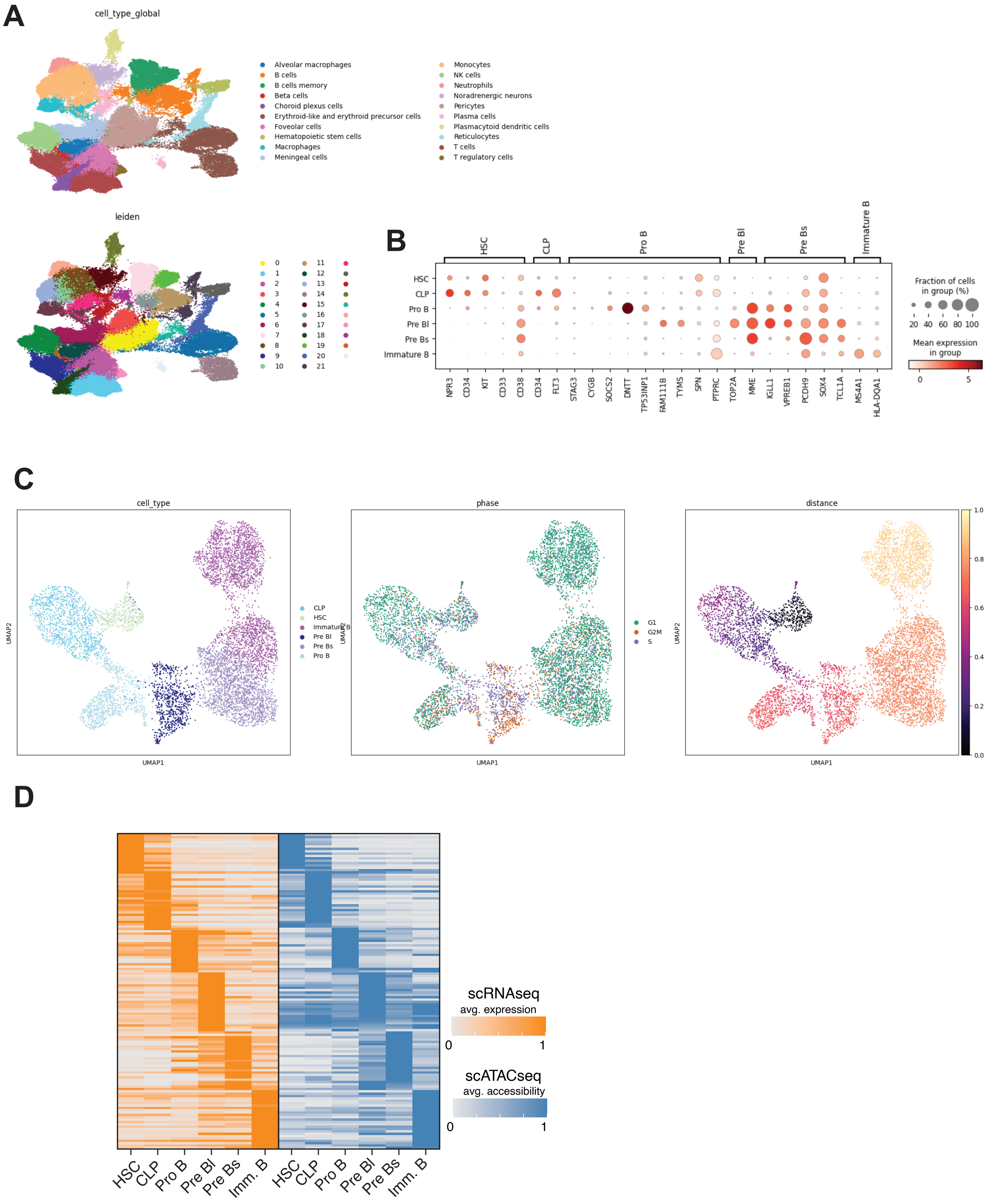
A
cell_type_global
Alveolar macrophages
B cells
B cells memory
Beta cells
Choroid plexus cells
Erythroid-like and erythroid precursor cells
Foveolar cells
Hematopoietic stem cells
Macrophages
Meningeal cells
Monocytes
NK cells
Neutrophils
Noradrenergic neurons
Pericytes
Plasma cells
Plasmacytoid dendritic cells
Reticulocytes
T cells
T regulatory cells
leiden
0 1 2 3 4 5 6 7 8 9 10 11 12 13 14 15 16 17 18 19 20 21
B
HSC
CLP
Pro B
Pre BI
Pre Bs
Immature B
NPR3 CD34 KIT CD33 CD38 CD34 FLT3 STAG3 CYGB SOCS2 DNTT TP53INP1 FAM111B TYMS SPN PTPRC TOP2A MME IGLL1 VPREB1 PCDH9 SOX4 TCL1A MS4A1 HLA-DQA1
Fraction of cells in group (%)
20 40 60 80 100
Mean expression in group
0 5
C
cell_type
CLP
HSC
Immature B
Pre BI
Pre Bs
Pro B
UMAP1
UMAP2
phase
G1
G2M
S
distance
0.0 0.2 0.4 0.6 0.8 1.0
D
scRNAseq
avg. expression
0 1
scATACseq
avg. accessibility
0 1
HSC CLP Pro B Pre BI Pre Bs Imm. B HSC CLP Pro B Pre BI Pre Bs Imm. B

**Figure S2. Characterization of B-cell precursor subsets from healthy donor bone marrow multiome data.**

**A)** UMAPs of scRNA-seq profiles from 10x Genomics Multiome data integrated across 10 healthy bone marrow donors. Top: global clustering of all hematopoietic populations with cell type annotations. Bottom: donor-specific sample identifiers.

**B)** Dot plot showing expression of canonical marker genes across B-cell precursor stages (HSC, CLP, Pro-B, Pre-BI, Pre-BII, Pre-Bs, Immature B). Dot size indicates the fraction of cells expressing each gene and color scale indicates mean expression.

**C)** UMAP embeddings of B-cell precursor subsets colored by cell type (left), cell cycle phase (middle), or diffusion pseudotime distance (right).

**D)** Heatmap of average scRNA-seq expression (orange) and scATAC-seq chromatin accessibility (blue) for stage-specific genes across B-cell precursor stages (HSC through Immature B).

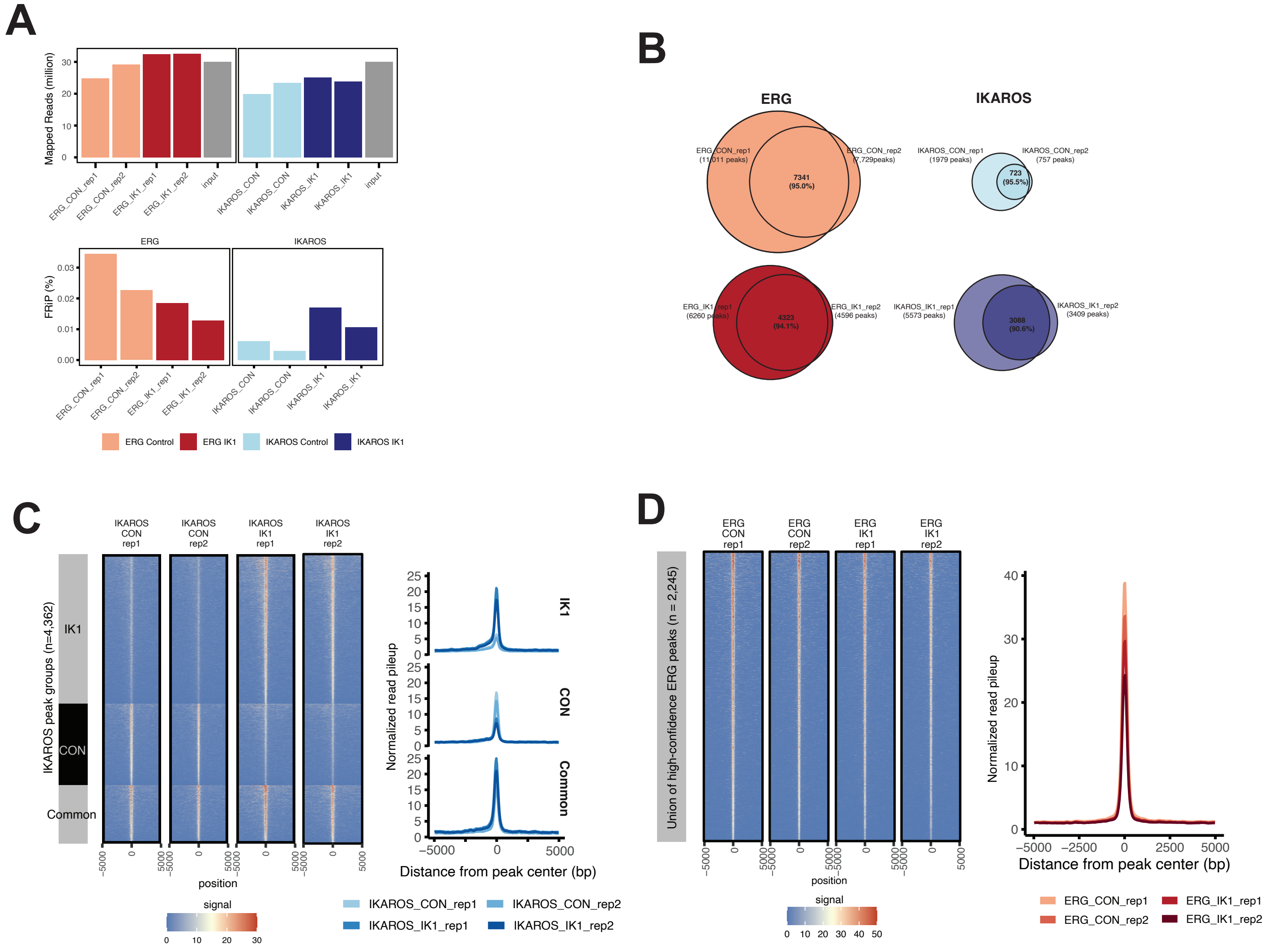
A
Mapped Reads (million)
ERG_CON_rep1
ERG_CON_rep2
ERG_IK1_rep1
ERG_IK1_rep2
input
IKAROS_CON
IKAROS_CON
IKAROS_IK1
IKAROS_IK1
input
ERG
IKAROS
FRIP (%)
ERG Control
ERG IK1
IKAROS Control
IKAROS IK1
B
ERG
IKAROS
ERG_CON_rep1 (11,011 peaks)
ERG_CON_rep2 (7,729peaks)
7341 (95.0%)
IKAROS_CON_rep1 (1979 peaks)
IKAROS_CON_rep2 (757 peaks)
723 (95.5%)
ERG_IK1_rep1 (6260 peaks)
ERG_IK1_rep2 (4596 peaks)
4323 (94.1%)
IKAROS_IK1_rep1 (5573 peaks)
IKAROS_IK1_rep2 (3409 peaks)
3088 (90.6%)
C
IKAROS CON rep1
IKAROS CON rep2
IKAROS IK1 rep1
IKAROS IK1 rep2
IKAROS peak groups (n=4,362)
IK1
CON
Common
position
signal
Normalized read pileup
Distance from peak center (bp)
IKAROS_CON_rep1
IKAROS_CON_rep2
IKAROS_IK1_rep1
IKAROS_IK1_rep2
D
ERG CON rep1
ERG CON rep2
ERG IK1 rep1
ERG IK1 rep2
Union of high-confidence ERG peaks (n = 2,245)
position
signal
Normalized read pileup
Distance from peak center (bp)
ERG_CON_rep1
ERG_CON_rep2
ERG_IK1_rep1
ERG_IK1_rep2

**Figure S3. Quality control for ERG and IKAROS ChIP-seq datasets across replicates.**

**A)** Sequencing depth and fraction of reads in peaks (FRiP percentage) for ERG and IKAROS ChIP-seq libraries. Top: Bar plots showing the number of mapped reads (in millions) for each replicate and input sample. Bottom: Fraction of Reads in Peaks (FRiP) percentages calculated using MACS2-called peaks for each replicate independently without IDR (Irreproducible Discovery Rate) filtering.

**B)** Venn diagrams showing peak overlap between biological replicates. Peak overlaps for ERG (left) and IKAROS (right) in both CON and IK1 conditions assessed using MACS2-called peaks for each replicate. Peaks were considered overlapping if they shared ≥1 bp with a max gap of 100 bp. These comparisons were performed prior to any IDR-based filtering for an inclusive view of replicate similarity.

**C, D)** ChIP-seq signal visualization over peak sets. Left: Heatmaps for RPKM-normalized ChIP-seq signal for IKAROS across distinct peak categories (IK1, CON, or shared). Aggregated line plots of ChIP-seq signal centered on the same peak sets, separated by condition and replicate. Right: Heatmaps for ERG ChIP-seq signal from both CON and IK1 ERG replicates centered on the union of IDR-filtered high-confidence ERG peaks across conditions.

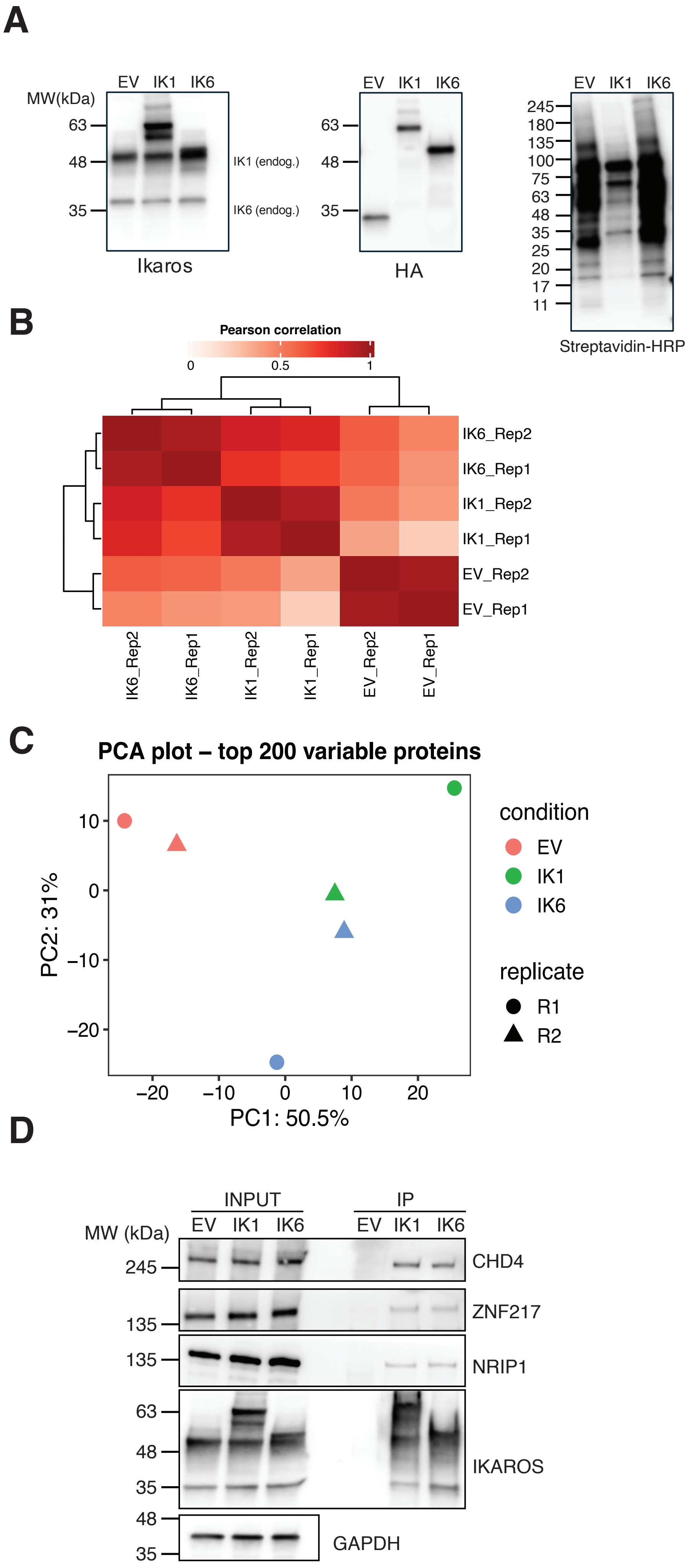

A
EV IK1 IK6
MW(kDa)
63
48
35
IK1 (endog.)
IK6 (endog.)
Ikaros
HA
245
180
135
100
75
25
20
17
11
Streptavidin-HRP
B
Pearson correlation
0
0.5
1
IK6_Rep2
IK6_Rep1
IK1_Rep2
IK1_Rep1
EV_Rep2
EV_Rep1
C
PCA plot – top 200 variable proteins
condition
EV
IK1
IK6
replicate
R1
R2
PC2: 31%
PC1: 50.5%
D
INPUT
IP
MW (kDa)
CHD4
ZNF217
NRIP1
IKAROS
GAPDH

**Figure S4. TurboID proximity labeling for IKAROS interactome profiling.**

**A)** Western blot validation of TurboID constructs expressed in PDX2 cells. Expression of endogenous IKAROS isoforms (IK1, IK6) and TurboID fusions was detected with IKAROS, HA, and streptavidin-HRP antibodies as indicated.

**B)** Pearson correlation heatmap of TurboID proteomics datasets showing high reproducibility across biological replicates and clustering by condition.

**C)** Principal component analysis (PCA) of the top 200 most variable proteins, based on DEP-normalized intensities, demonstrating clear separation of IK1, IK6, and empty vector (EV) samples across replicates.

**D)** Co-immunoprecipitation of IKAROS-associated proteins identified by TurboID. Whole-cell lysates from PDX2 cells expressing TurboID-empty vector (EV), TurboID-IK1, or TurboID-IK6 were immunoprecipitated using anti-HA–conjugated beads to capture HA-tagged TurboID fusion proteins and immunoblotted for CHD4, ZNF217, NRIP1, and IKAROS. Molecular-weight markers (kDa) are shown on the left. GAPDH serves as a loading control for input samples.

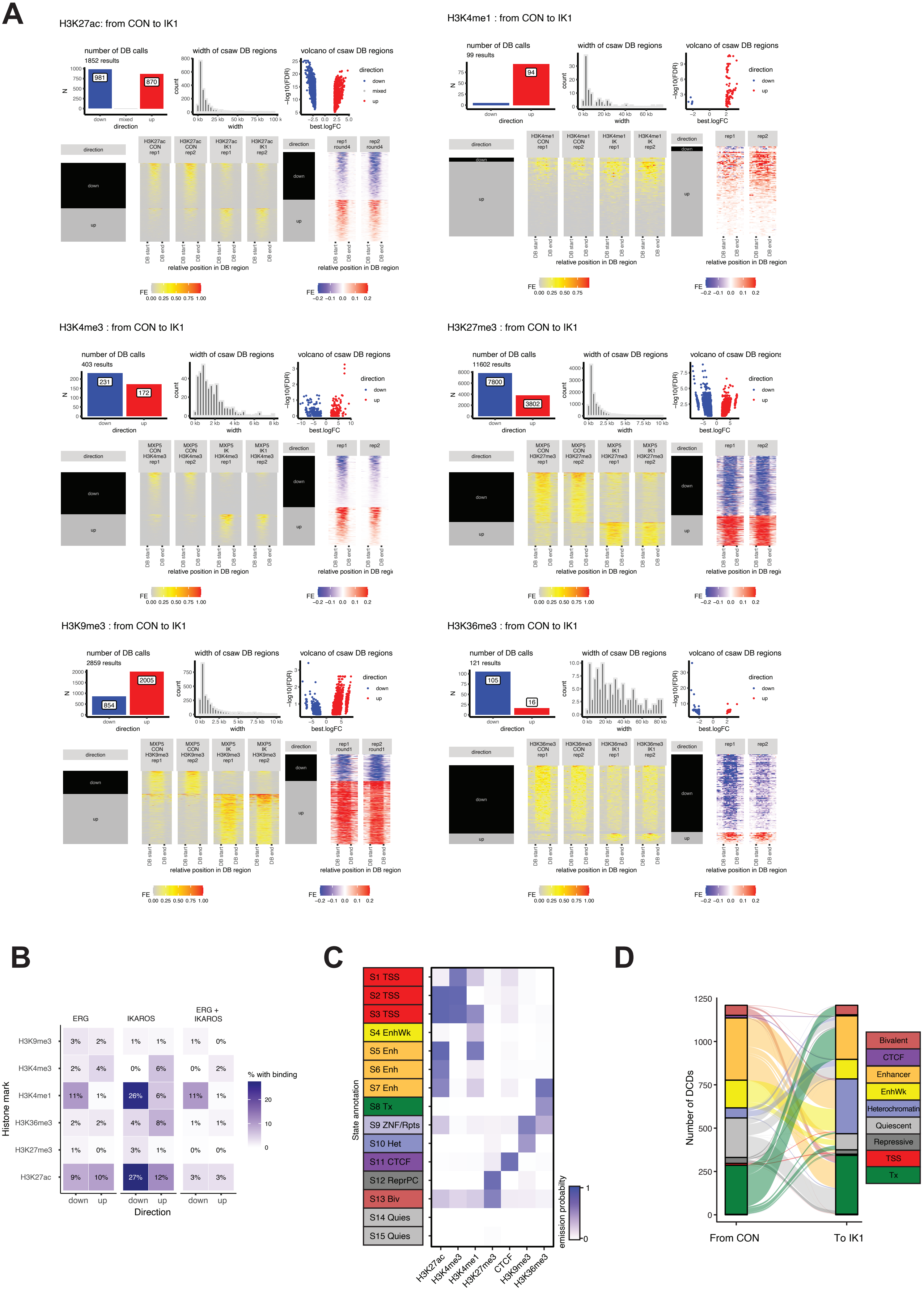

A
H3K27ac: from CON to IK1
number of DB calls
1852 results
981
870
width of csaw DB regions
volcano of csaw DB regions
direction
down
mixed
up
best.logFC
relative position in DB region
H3K4me1 : from CON to IK1
99 results
94
H3K4me3 : from CON to IK1
403 results
231
172
H3K27me3 : from CON to IK1
11602 results
7800
3802
H3K9me3 : from CON to IK1
2859 results
854
2005
H3K36me3 : from CON to IK1
121 results
105
16
FE
B
ERG
IKAROS
ERG + IKAROS
H3K9me3
H3K4me3
H3K4me1
H3K36me3
H3K27me3
H3K27ac
Histone mark
% with binding
Direction
C
S1 TSS
S2 TSS
S3 TSS
S4 EnhWk
S5 Enh
S6 Enh
S7 Enh
S8 Tx
S9 ZNF/Rpts
S10 Het
S11 CTCF
S12 ReprPC
S13 Biv
S14 Quies
S15 Quies
State annotation
emission probability
H3K27ac
H3K4me3
H3K4me1
H3K27me3
CTCF
H3K9me3
H3K36me3
D
Number of DCDs
From CON
To IK1
Bivalent
CTCF
Enhancer
EnhWk
Heterochromatin
Quiescent
Repressive
TSS
Tx

**Figure S5. Chromatin state and histone modification changes following IK1 induction in MXP5 cells.**

**A)** Validation of differential binding (DB) calls from csaw for six histone marks (H3K27ac, H3K4me1, H3K4me3, H3K27me3, H3K9me3, H3K36me3). For each mark, summary plots show the number of significant DB regions (FDR < 0.05), width distributions, and volcano plots of fold change versus mean abundance. Heatmaps display normalized read density (±2 kb) across DB regions, with separate tracks for up- and downregulated peaks.

**B)** Overlap between DB regions for each histone modification and transcription factor binding. Percentages indicate the fraction of DB regions intersecting ERG, IKAROS, or shared ERG + IKAROS ChIP-seq peaks.

**C)** ChromHMM was used to segment the genome into 15 chromatin states in MXP5 cells based on combinatorial patterns of six histone modifications (H3K27ac, H3K4me1, H3K4me3, H3K27me3, H3K9me3, H3K36me3) and CTCF occupancy. Condition- and replicate-specific BAM files were binarized at 200 bp resolution with matched input controls, and states were learned jointly across control and IK1-induced conditions. The heatmap displays the emission probabilities for each mark across the 15 learned chromatin states. States were annotated as follows: active promoters (S1–S3), weak and strong enhancers (S4–S7), transcriptional (S8), ZNF/repeats (S9), heterochromatin (S10), CTCF-bound regions (S11), repressive Polycomb (S12), bivalent (S13), and quiescent or low-signal states (S14–S15).

**D)** Dynamic chromatin domains (DCDs) were identified using SCIDDO [93], which compares chromatin state segmentations between conditions to detect significant state transitions. ChromHMM segmentations from MXP5 cells under CON and IK1-induced conditions were generated using a 15-state model and used as input for SCIDDO. DCDs were defined as genomic intervals with significant state transitions between conditions, based on the most probable ChromHMM state in each condition. Transition frequencies were quantified and visualized using Sankey plots to highlight genome-wide shifts in chromatin state. State annotations were grouped into promoter-associated (S1–S3), enhancer (S4–S7), transcriptional (S8), repressive Polycomb (S12), bivalent (S13), heterochromatin (S9, S10), CTCF-bound (S11), and quiescent (S14–S15) categories. Enhancer- and transcription-associated states (S4–S8) exhibited the largest number of transitions between conditions, while promoter and heterochromatin states showed fewer changes overall.

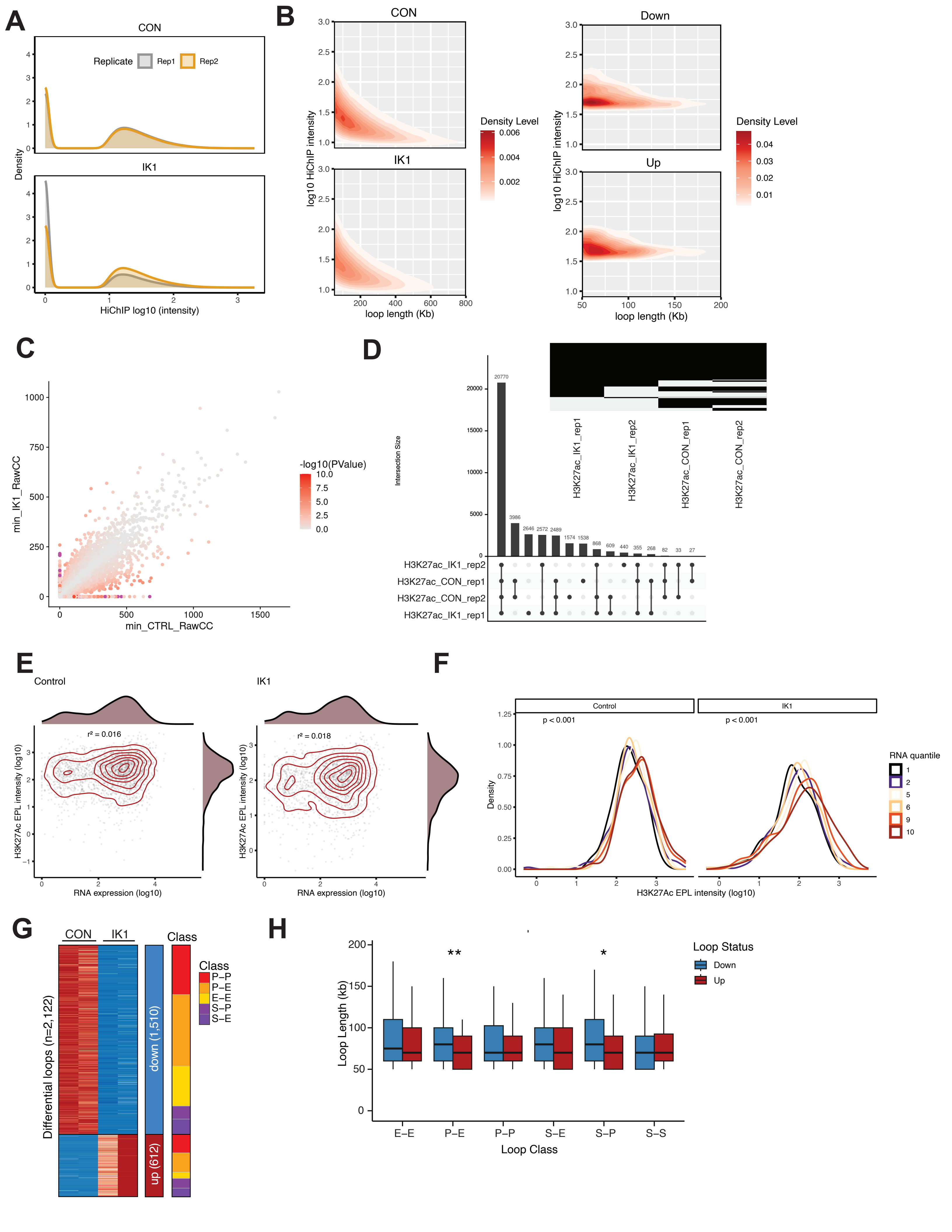

A
CON
Replicate
Rep1
Rep2
Density
IK1
HiChIP log10 (intensity)
B
CON
IK1
log10 HiChIP intensity
loop length (Kb)
Density Level
Down
Up
C
min_IK1_RawCC
min_CTRL_RawCC
-log10(PValue)
D
Intersection Size
H3K27ac_IK1_rep1
H3K27ac_IK1_rep2
H3K27ac_CON_rep1
H3K27ac_CON_rep2
E
Control
IK1
r² = 0.016
r² = 0.018
H3K27Ac EPL intensity (log10)
RNA expression (log10)
F
Control
IK1
p < 0.001
RNA quantile
Density
H3K27Ac EPL intensity (log10)
G
CON
IK1
Class
P–P
P–E
E–E
S–P
S–E
Differential loops (n=2,122)
down (1,510)
up (612)
H
Loop Status
Down
Up
Loop Length (kb)
Loop Class
E–E
P–E
P–P
S–E
S–P
S–S

**Figure S6. Differential H3K27ac HiChIP loops and UMI-4C in MXP5 cells.**

**A)** Heatmap of chromatin loops with significant differential contact frequency ($padj < 0.05$, $|\log_2$ fold-change$| > 1$) between control (CON) and IK1 conditions across biological replicates ($n = 2$ per condition). Loops are stratified by loop class: promoter–promoter (P–P), promoter–enhancer (P–E), enhancer–enhancer (E–E), enhancer–structural (E–S), structural–enhancer (S–E), structural–promoter (S–P), and structural–structural (S–S), where S indicates overlap with structural CTCF anchors. Loops with increased contact frequency in IK1 are shown in red and decreased loops in blue.

**B)** Boxplots of loop lengths for up- and down-regulated loops across classes. Differences in loop length distributions were assessed by Wilcoxon rank-sum test (*$p < 0.05$, **$p < 0.01$).

**C)** Quality control metrics for CCND2 UMI-4C libraries. Stacked barplots show read specificity, filtering, and alignment categories for each replicate (n = 2 CON, n = 2 IK1). Total UMI counts per library are shown below. The UMI-4C domainogram displays interaction frequencies across the CCND2 locus, with color intensity reflecting local signal enrichment.

**D)** Chromatin architecture and regulatory features at the *CCND2* locus. H3K27ac HiChIP contact maps for CON and IK1 are shown with differential loops (gained in red, lost in blue). Genome browser tracks display ChIP-seq profiles (H3K27ac, IKAROS, ERG, CTCF), bulk ATAC-seq, pseudobulk scATAC-seq, and RNA-seq in CON and IK1. UMI-4C contact profiles from the *CCND2* bait are plotted below, with the y-axis showing the smoothed UMI-4C trend signal.

**E)** Immunoblot of IKAROS and CCND2 protein in MXP5 cells at indicated timepoints (hours) after DOX-induced IK1 expression. GAPDH is shown as a loading control.

**F)** UMI-4C quantitative analysis of chromatin contacts in MXP5 cells under control and IK1 conditions. Baits are designed against *CCND2,* two independent *MYC* bait positions and *CCDC26*. On the left are quality control metrics for UMI-4C libraries for the indicated baits. Stacked barplots show the distribution of read specificity, filtering, and alignment categories for each replicate (n = 2 CON, n = 2 IK1 per bait), and a summary of the number of unique UMIs obtained per library. Contact profiles (right) display normalized UMI-4C trend signal across the indicated genomic loci (chr12: 3.9-4.5 Mb for *CCND2*; chr8:127–129 Mb for *MYC*; chr8:129–130.5 Mb for *CCDC26*). Arrowheads mark bait positions, and annotated genes are shown above each track. Color scales indicate $\log_2$ odds ratio (OR) and $\log_2$ fold-change (FC) in UMI counts across replicates.

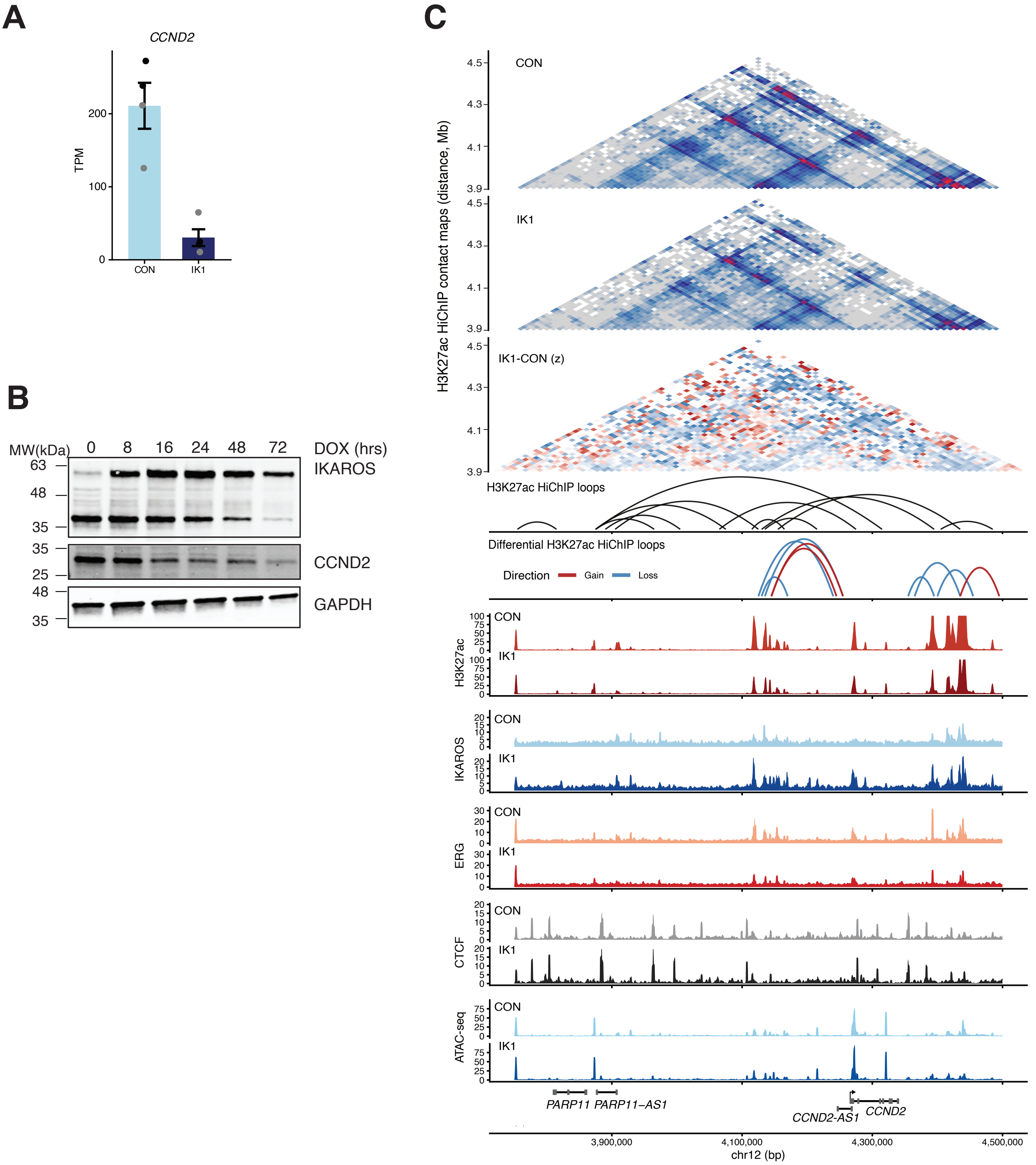

A
CCND2
TPM
200
100
0
CON
IK1
B
MW(kDa)
0
8
16
24
48
72
DOX (hrs)
63
48
35
IKAROS
35
25
CCND2
48
35
GAPDH
C
H3K27ac HiChIP contact maps (distance, Mb)
CON
IK1
IK1-CON (z)
4.5
4.3
4.1
3.9
H3K27ac HiChIP loops
Differential H3K27ac HiChIP loops
Direction
Gain
Loss
H3K27ac
IKAROS
ERG
CTCF
ATAC-seq
CON
IK1
PARP11
PARP11-AS1
CCND2-AS1
CCND2
3,900,000
4,100,000
4,300,000
4,500,000
chr12 (bp)

D

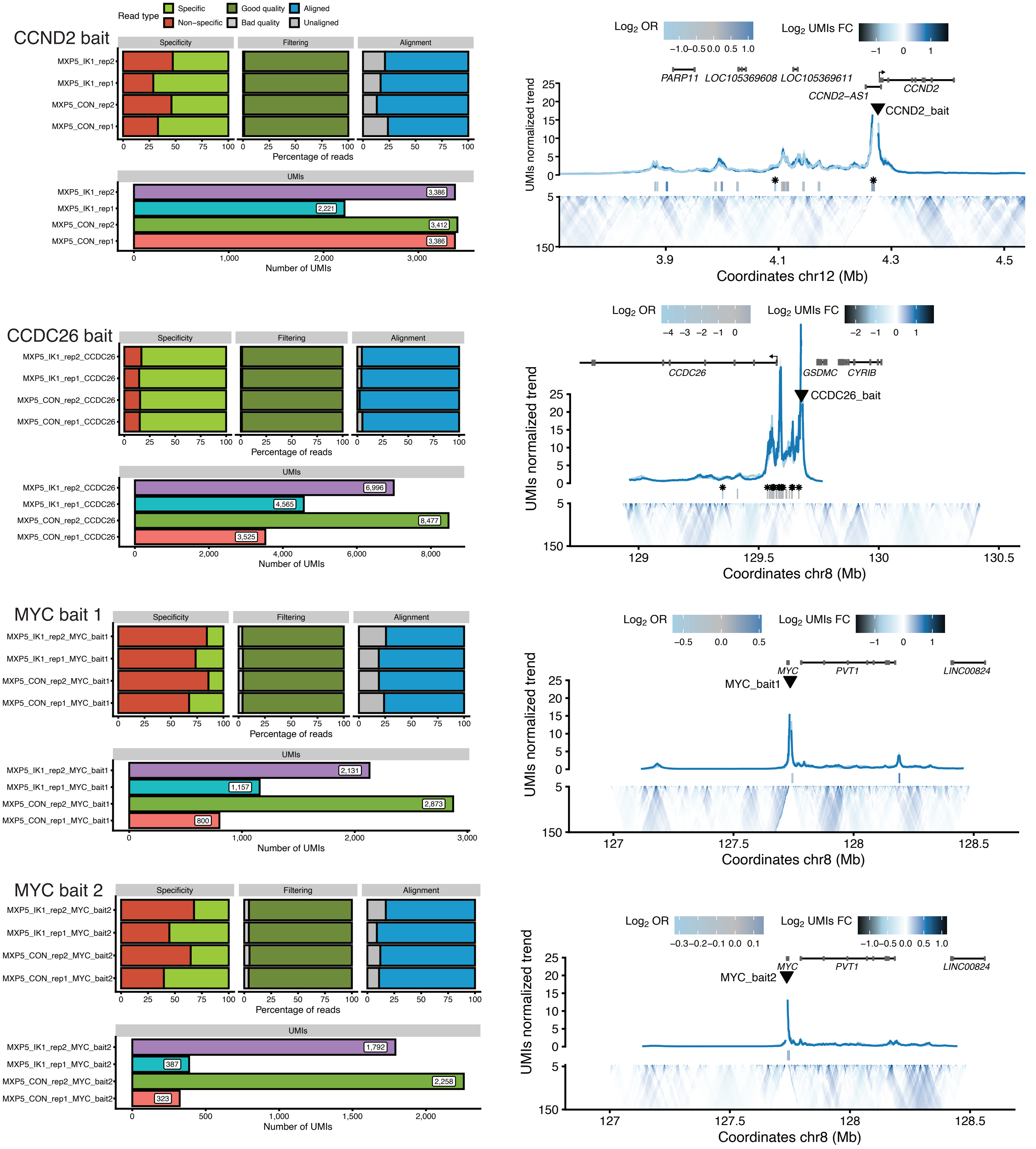

Read type
Specific
Non−specific
Good quality
Bad quality
Aligned
Unaligned
CCND2 bait
Specificity
Filtering
Alignment
MXP5_IK1_rep2
MXP5_IK1_rep1
MXP5_CON_rep2
MXP5_CON_rep1
Percentage of reads
UMIs
3,386
2,221
3,412
3,386
Number of UMIs
Log2 OR
Log2 UMIs FC
PARP11
LOC105369608
LOC105369611
CCND2−AS1
CCND2
CCND2_bait
UMIs normalized trend
Coordinates chr12 (Mb)
CCDC26 bait
MXP5_IK1_rep2_CCDC26
MXP5_IK1_rep1_CCDC26
MXP5_CON_rep2_CCDC26
MXP5_CON_rep1_CCDC26
6,996
4,565
8,477
3,525
CCDC26
GSDMC
CYRIB
CCDC26_bait
Coordinates chr8 (Mb)
MYC bait 1
MXP5_IK1_rep2_MYC_bait1
MXP5_IK1_rep1_MYC_bait1
MXP5_CON_rep2_MYC_bait1
MXP5_CON_rep1_MYC_bait1
2,131
1,157
2,873
800
MYC
PVT1
LINC00824
MYC_bait1
MYC bait 2
MXP5_IK1_rep2_MYC_bait2
MXP5_IK1_rep1_MYC_bait2
MXP5_CON_rep2_MYC_bait2
MXP5_CON_rep1_MYC_bait2
1,792
387
2,258
323
MYC_bait2

**Figure S7. CCND2 regulation and chromatin interactions at target loci in CON and IK1 conditions.**

**A)** CCND2 mRNA expression in MXP5 and PDX2 cells under control (CON) and IK1 conditions. RNA-seq TPM values are shown for two biological replicates per condition ($n = 2$). Black dots = MXP5; grey dots = PDX2. Bars indicate mean ± SD.

**B)** Western blot analysis of IKAROS and CCND2 protein expression in MXP5 cells. Doxycycline (DOX) was added at time 0 to induce IK1 expression. Lysates were collected at the indicated timepoints and probed for IKAROS, CCND2, and GAPDH (loading control).

**C)** Chromatin architecture and regulatory landscape at the *CCND2* locus (chr12:3.9–4.5 Mb). Top: H3K27ac HiChIP contact matrices in control and IK1 conditions, along with a differential Z-score matrix (IK1 − CON) showing relative changes in contact intensity. Middle: Significant H3K27ac HiChIP loops are shown as arcs. Black arcs represent significant H3K27ac HiChIP loops supported in both CON and IK1 replicates ($q \leq 0.01$ in each), and anchored within ±40 kb of the CCND2 promoter. Colored arcs represent differential loops (IK1 vs CONl;$|\log_2 FC| \geq 0.5$, $P < 0.05$, replicate support), with gains in red and losses in blue. Bottom: ChIP-seq signals for H3K27ac, IKAROS, ERG, and CTCF, and bulk ATAC-seq in CON and IK1 MXP5 cells. Tracks are RPKM-normalized, with condition-specific color coding.

**D)** UMI-4C quality control and contact profiles for targeted loci (*CCND2*, *CCDC26*, and *MYC*). Left: Library quality metrics for each bait and replicate (n = 2 CON, n = 2 IK1 per bait), showing read specificity, filtering categories, alignment percentages, and total unique UMI counts. Right: Normalized UMI-4C contact profiles for each bait. Interaction frequencies are plotted along the genomic axis. Vertical colored bars indicate peak interaction signals (light blue: CON; dark blue: IK1; gray: shared). Asterisks denote significant differential interactions ($FDR < 0.05$).

Arrowheads mark the bait positions. Color scales above each track represent $log_2$ odds ratio (OR) and $log_2$ fold change (FC) in UMI counts between conditions.

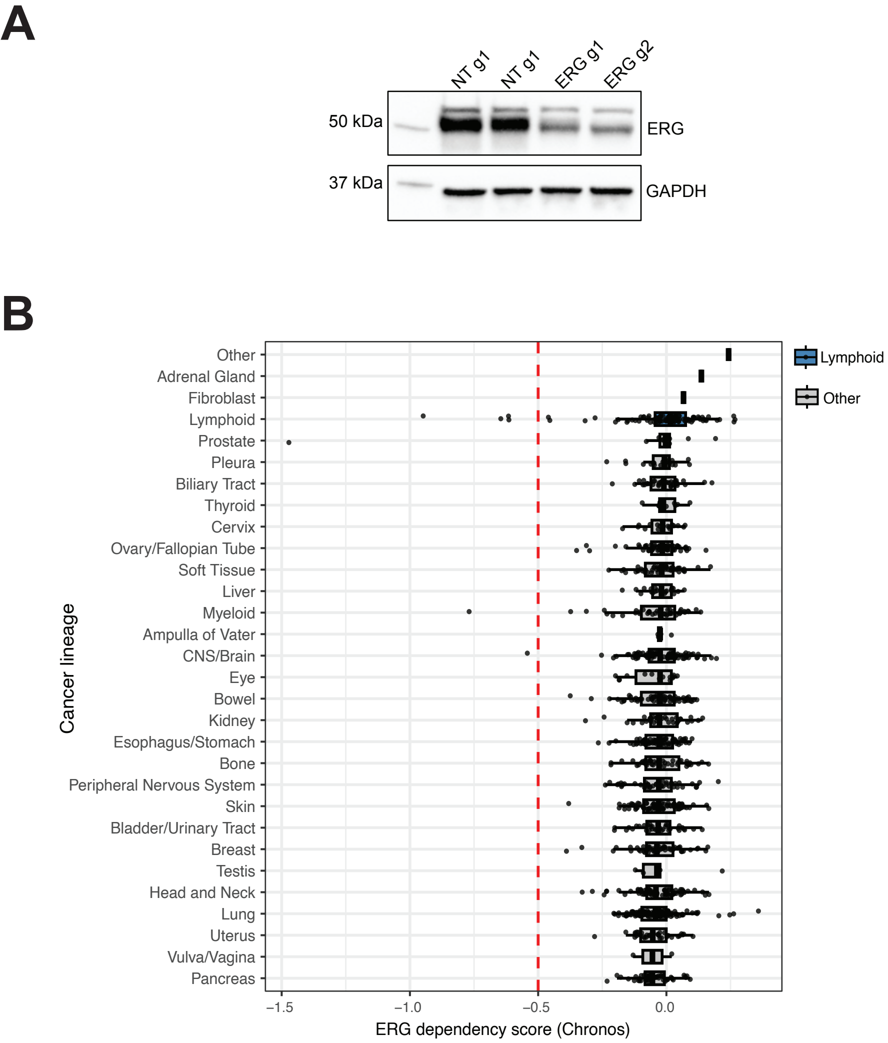
A
NT g1
NT g1
ERG g1
ERG g2
50 kDa
ERG
37 kDa
GAPDH
B
Other
Adrenal Gland
Fibroblast
Lymphoid
Prostate
Pleura
Biliary Tract
Thyroid
Cervix
Ovary/Fallopian Tube
Soft Tissue
Liver
Myeloid
Ampulla of Vater
CNS/Brain
Eye
Bowel
Kidney
Esophagus/Stomach
Bone
Peripheral Nervous System
Skin
Bladder/Urinary Tract
Breast
Testis
Head and Neck
Lung
Uterus
Vulva/Vagina
Pancreas
Cancer lineage
–1.5
–1.0
–0.5
0.0
ERG dependency score (Chronos)
Lymphoid
Other

**Figure S8. ERG depletion and lineage-specific dependency in B-ALL and other cancer cell lines.**

**A)** Western blot confirming ERG protein knockdown in PDX2 cells following CRISPR interference (CRISPRi) targeting the ERG promoter region (TSS) using two independent sgRNAs (ERG g1 and ERG g2). Non-targeting sgRNAs (NT g1) serve as controls. GAPDH is shown as a loading control.

**B)** ERG dependency scores across 1,000+ cancer cell lines from the Cancer Dependency Map (DepMap, 22Q4), grouped by lineage. Each point represents a cell line; boxplots indicate median and interquartile range. Lymphoid cell lines (blue) showed lower Chronos scores (greater dependency) compared to other lineages (gray). Dashed red line indicates the Chronos dependency threshold (–0.5). The lymphoid subset is shown separately in Fig. 7F.
