## Supplemental Table Legends for "IKAROS Gene Regulatory Network Reveal ERG as a Vulnerability in B-cell Acute Lymphoblastic Leukemia"

**Table S1. RNA-seq differential expression analysis results comparing IK1 vs control in MXP5 and PDX2 cells.** Normalized gene expression differences between IK1-induced and control cells, calculated using DESeq2, including gene names, baseMean, log₂ fold change, standard error, Wald statistic, p-value, adjusted p-value (padj).

**Table S2. Chromatin accessibility changes from differential accessibility analysis comparing IK1 vs control in MXP5 and PDX2 cells.** ATAC-seq peaks showing differential accessibility between IK1-induced and control MXP5 and PDX2 cells. Includes all tested regions with genomic coordinates, log₂ fold change, adjusted p-values, and classification as more accessible (open), less accessible (closed), or not significant (ns).

**Table S3. Gene Set Enrichment Analysis from differential expression analysis results comparing IK1 vs control in MXP5 and PDX2 cells.** Enriched pathways from GSEA using MSigDB Hallmark and Canonical Pathways (C2:CP) gene sets. Gene set name, normalized enrichment score (NES), nominal p-value, FDR q-value, and leading-edge genes.

**Table S4. Upstream regulator analysis from differential expression analysis results comparing IK1 vs control in MXP5 and PDX2 cells.** Predicted transcriptional regulators based on differentially expressed genes (IK1 vs Control) generated using Ingenuity Pathway Analysis.

**Table S5.** **Transcription factor footprinting results**. Differential TF binding site accessibility inferred from TOBIAS footprinting analysis of ATAC-seq profiles. TF name, footprint score changes, p-values, and binding direction.

**Table S6.** **Gene regulatory network inference.** Condition-specific TF–target regulatory relationships predicted from integrated RNA-seq and ATAC-seq data using PECA. TF, target gene, edge weight, correlation, activity, fold change, and condition.

**Table S7.** **IKAROS and ERG ChIP-seq consensus peaks.** Consensus peaks derived from replicate analysis of ChIP-seq data for IKAROS and ERG under control and IK1-induced conditions. Genomic coordinates, peak identifier, treatment group, score, signalValue, pValue, qValue, and peak summit position.

**Table S8. IKAROS proximity interactome results.** Proteins significantly enriched in TurboID-IKZF1 or TurboID-IKZF6 versus TurboID-NLS control samples. UniProt IDs, gene symbols, log₂ fold change, adjusted p-values, and enrichment results from GO Molecular Function and Reactome 2022 pathway analyses.

**Table S9. Genome-wide differential chromatin loops identified by H3K27ac HiChIP following IK1 induction.** High-confidence loops significantly altered (FDR < 0.05) between control and IK1-induced MXP5 cells, annotated with genomic coordinates, statistical values (logFC, logCPM, p-value, FDR), total interaction counts, and the direction of change (gain/loss).

**Table S10. UMI-4C oligonucleotides, CRISPRi sgRNA sequences, antibodies and cell lines.**
All oligonucleotide sequences used in the study, including primers and indexed Illumina-compatible adapters for UMI-4C library preparation (bait primers, downstream primers, and barcode-adapter sequences for multiplexed sequencing), CRISPRi sgRNA sequences, antibody information and cell lines.
